# An end-to-end workflow for non-destructive 3D pathology

**DOI:** 10.1101/2023.08.03.551845

**Authors:** Kevin W. Bishop, Lindsey A. Erion Barner, Qinghua Han, Elena Baraznenok, Lydia Lan, Chetan Poudel, Gan Gao, Robert B. Serafin, Sarah S.L. Chow, Adam K. Glaser, Andrew Janowczyk, David Brenes, Hongyi Huang, Dominie Miyasato, Lawrence D. True, Soyoung Kang, Joshua C. Vaughan, Jonathan T.C. Liu

## Abstract

Recent advances in 3D pathology offer the ability to image orders-of-magnitude more tissue than conventional pathology while providing a volumetric context that is lacking with 2D tissue sections, all without requiring destructive tissue sectioning. Generating high-quality 3D pathology datasets on a consistent basis is non-trivial, requiring careful attention to many details regarding tissue preparation, imaging, and data/image processing in an iterative process. Here we provide an end-to-end protocol covering all aspects of a 3D pathology workflow (using light-sheet microscopy as an illustrative imaging platform) with sufficient detail to perform well-controlled preclinical and clinical studies. While 3D pathology is compatible with diverse staining protocols and computationally generated color palettes for visual analysis, this protocol will focus on a fluorescent analog of hematoxylin and eosin (H&E), which remains the most common stain for gold-standard diagnostic determinations. We present our guidelines for a broad range of end-users (e.g., biologists, clinical researchers, and engineers) in a simple tutorial format.

## Introduction

Anatomic pathology, which entails the microscopic analysis of biological tissues, is often regarded as the gold-standard for clinical diagnostics/prognostics and for many preclinical studies. While conventional slide-based 2D histology remains the most widespread approach for high-resolution interrogation of tissue specimens, 3D pathology of intact tissues is gaining attention for both research applications and potential clinical assays^1,2^. Compared to conventional pathology, 3D pathology interrogates orders-of-magnitude more tissue and provides a feature-rich volumetric context for complex tissue morphologies, all without destructive tissue sectioning. This novel volumetric “big data” has the potential to improve diagnostic sensitivity and accuracy, prognostication of patient outcomes, and prediction of treatment response in comparison to 2D pathology, with the ultimate goal of improving clinical decisions for diverse individuals.

To accelerate the preclinical and clinical adoption of 3D pathology, well-controlled studies with large, curated sets of tissue specimens are needed. Here we present a comprehensive, end-to-end 3D pathology workflow termed Path3D to guide researchers in generating consistent, high-quality 3D-pathology datasets, with a focus on large-scale clinical and preclinical studies.

### Development of the protocol

A variety of approaches have been developed for staining, imaging, and visualizing pathology specimens in 3D with impressive proof-of-concept results^1,2^. However, consistently generating high-quality 3D pathology datasets across hundreds of samples for large-scale clinical studies is nontrivial, requiring refined protocols for tissue preparation, imaging, data processing, and quality control. Generating reliable data is complicated by the fact that no general microscopy end-user can be expected to be an expert in all stages of this cross-disciplinary pipeline. Therefore, a comprehensive and detailed protocol that can be implemented by general researchers would be of value. In addition to the technical workflow requirements (image resolution, dye selection, etc.), key practical requirements include quality control, throughput/ease-of-use for hundreds of specimens, and a visualization format that is familiar to researchers and clinicians. Here we attempt to address these requirements and to provide a drop-in alternative to conventional 2D histology. Our protocol is designed for formalin-fixed (FF) or formalin-fixed paraffin-embedded (FFPE) tissues, which describes the vast majority of archived clinical specimens. The final output of the workflow described here is a 3D dataset that can be viewed as a stack of 2D images by pathologists/researchers and that can optionally be false-colored to match the appearance of standard hematoxylin and eosin (H&E) histology to facilitate interpretation. While we do not discuss downstream image-processing and computational-analysis workflows, which will vary greatly between end-users, the H&E-analog 3D datasets generated by our protocol are also an ideal starting point for many of these computational efforts^3–5^.

Our Path3D protocol includes four main sections (Fig. 1): (1) tissue preparation (staining and optical clearing), (2) 3D microscopy, (3) data processing, and (4) quality control. The first three sections generate the initial datasets, which can mimic the appearance of standard H&E images if helpful (Fig. 2). The last section ensures high-quality results in large-scale studies and provides feedback for iterative parameter optimization within the first three sections, as is often needed during the early stages of a project. Path3D uses a relatively fast labeling and clearing method (a few days from start to finish) that can be reliably carried out by a student (including advanced undergraduates) or research technician. In summary, Path3D enables well-controlled clinical or preclinical studies where consistent, high-throughput 3D pathology datasets are needed for tens to hundreds of specimens.

**Fig. 1.**
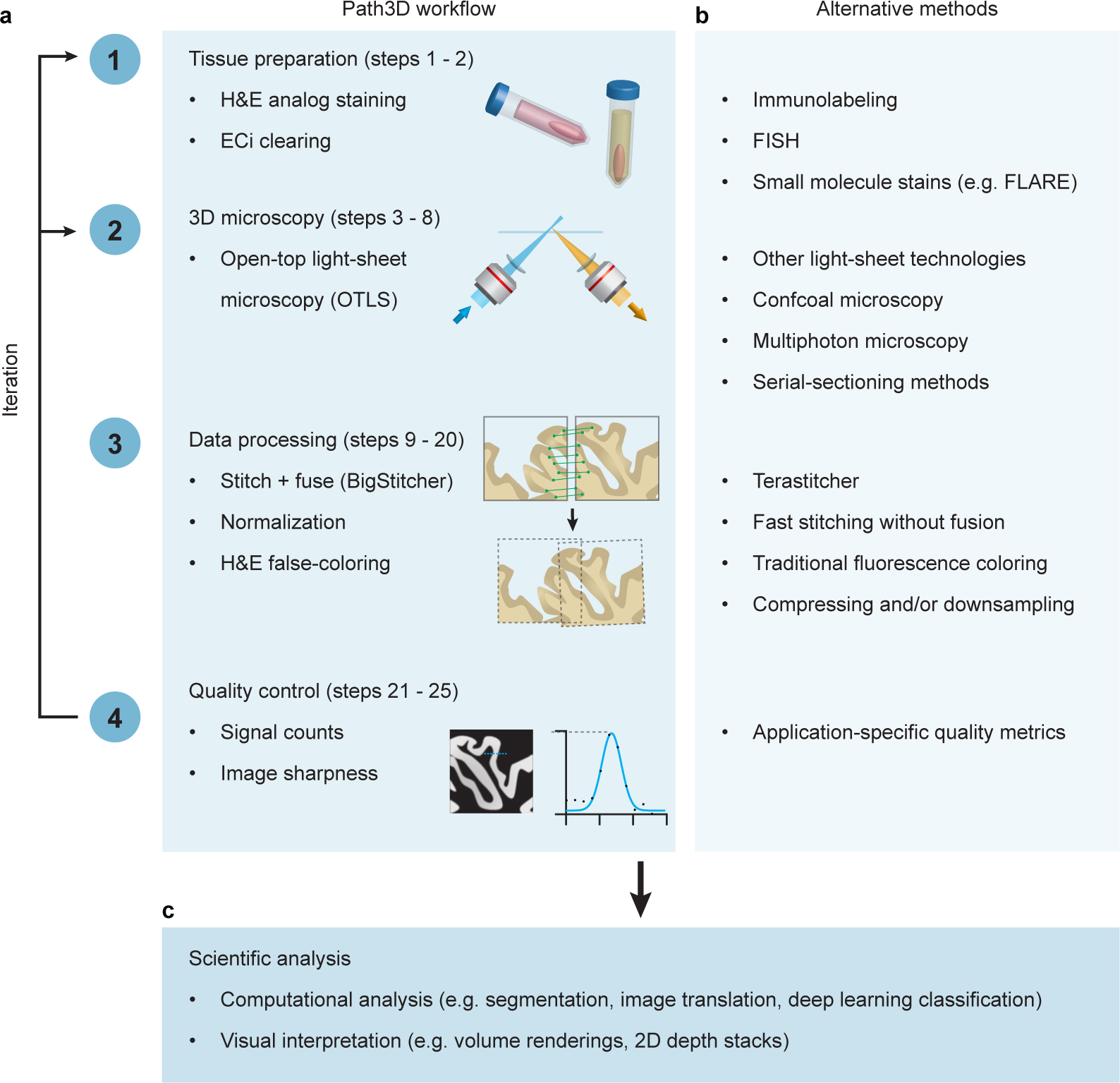
Protocol overview. **a**, Flowchart showing the four key sections to the Path3D workflow: tissue preparation, 3D microscopy, data processing, and quality control. Iterative optimization is essential for producing consistent and high-quality datasets. **b**, Each section of the standard Path3D protocol can be adapted based on the needs of a specific application. **c**, Path3D datasets are suitable for a range of scientific analyses based on computational analysis, visual interpretation, or a combination of the two.

**Fig. 2.**
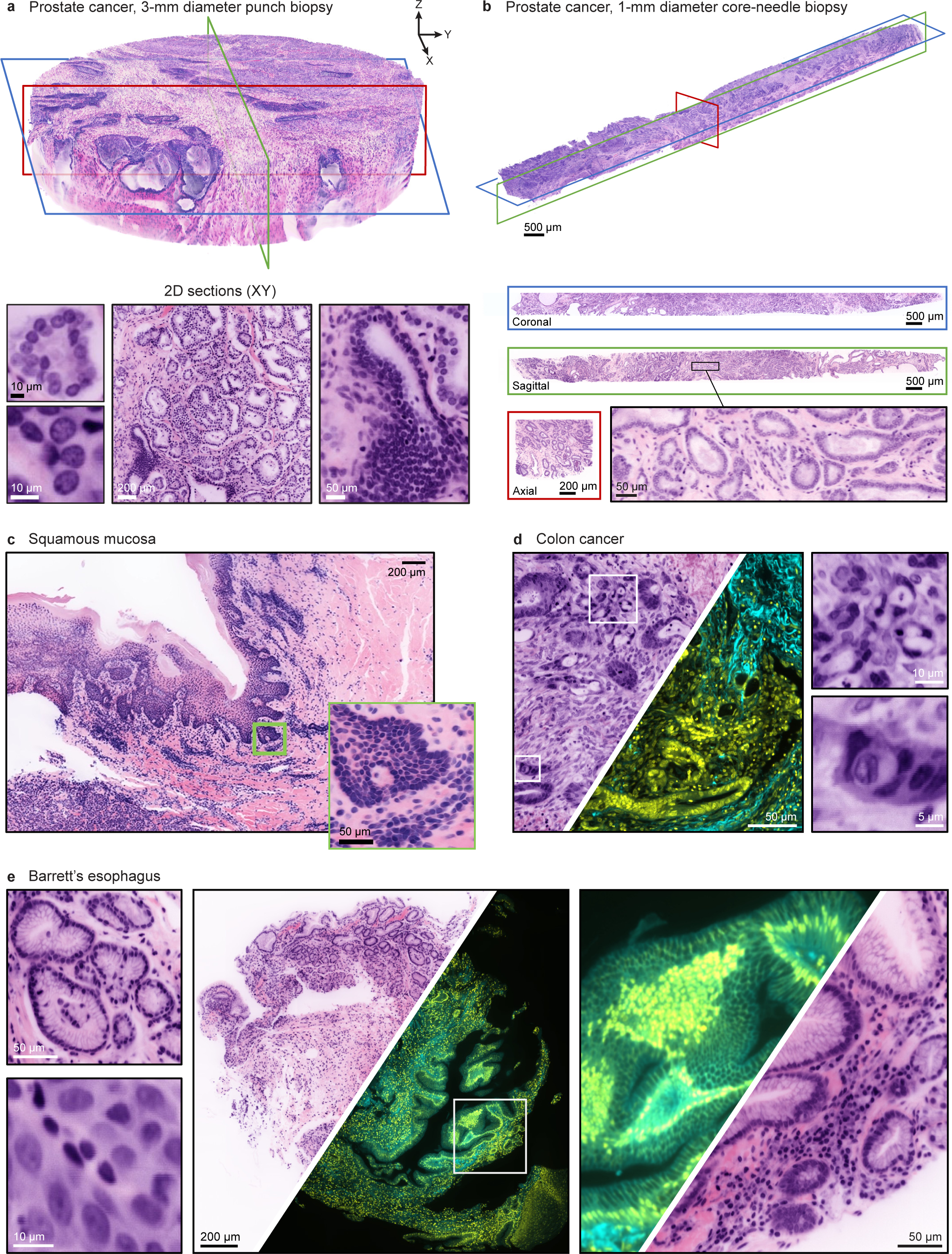
Image atlas showing Path3D datasets of archived human pathology specimens. **a**, 3D rendering (produced using Imaris) and *en face* 2D cross sections of a 3-mm diameter punch biopsy of prostate cancer tissue (FFPE) false colored to mimic the appearance of standard H&E staining. **b**, 3D rendering (produced using Imaris) and orthogonal 2D cross sections of a 1-mm diameter core-needle biopsy of prostate cancer tissue false colored to mimic the appearance of standard H&E staining. **c**, 2D cross section of squamous mucosal tissue false colored to mimic the appearance of standard H&E staining, reproduced from Serafin et al.^45^ (CC BY 4.0). **d**, 2D cross section of colon cancer tissue shown with both H&E false coloring and standard fluorescence coloring. **e**, 2D cross section of esophagus tissue (Barrett’s esophagus) shown with both H&E false coloring and standard fluorescence coloring. Images are shown with a standard sharpening filter. **! CAUTION** Experiments using human tissues must follow appropriate institutional and governmental regulations with respect to informed consent. The results in this figure were from de-identified tissues provided by an institutional tissue bank and are not considered human-subjects research.

### Protocol overview

The initial tissue-processing steps (Fig. 3) of Path3D are designed to mimic the traditional H&E stain while making the tissue transparent (optically clear) for high-throughput volumetric microscopy. The traditional absorption-based H&E stain for 2D histology offers general-purpose contrast by combining hematoxylin, which stains negatively charged DNA (i.e., nuclei) with a purple hue, and eosin, which stains positively charged proteins (i.e., cytoplasm and extracellular matrix) with a pink hue. To provide a fluorescent analog of standard H&E for 3D microscopy, Path3D likewise uses both a nuclear dye (TO-PRO-3) and a cytoplasmic dye (eosin, which is naturally fluorescent). The use of small molecule dyes is essential for efficient penetration into thick tissues. After tissue staining, optical clearing is used to render thick specimens optically transparent by homogenizing the refractive index within each sample, thereby enabling fluorescence imaging deep within the tissue. Path3D is optimized for tissue clearing with ethyl cinnamate (ECi)^6^, a solvent-based protocol similar to iDISCO methods^7,8^. Ethyl cinnamate clearing is relatively simple and uses nontoxic reagents.

**Fig. 3.**
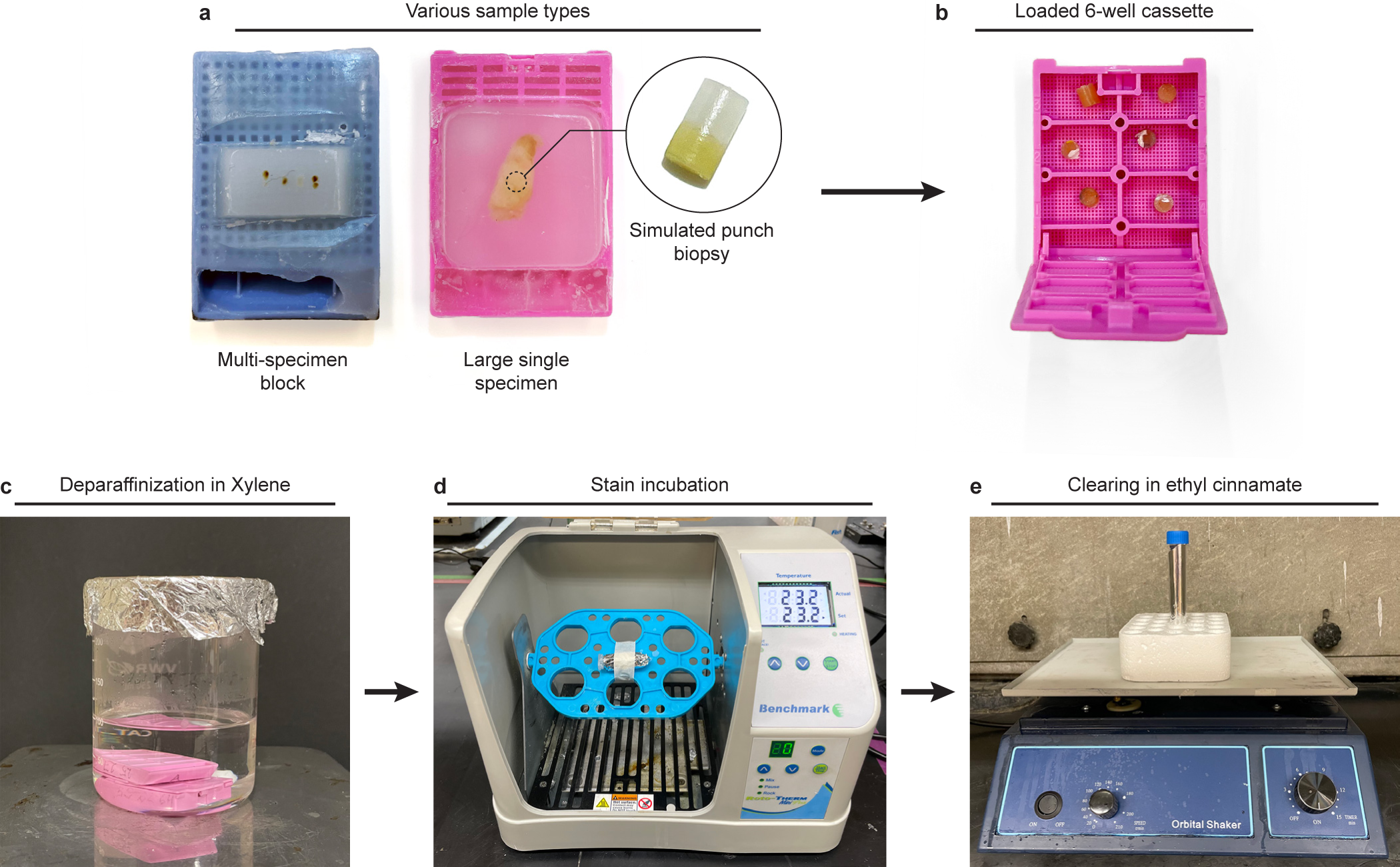
Photos showing key tissue processing steps. **a**, Various sample types can be used with Path3D including common archival specimens such as multi-specimen blocks, large single specimens, or punch biopsies. **b**, Tissue punches are loaded into 6-well cassettes for deparaffinization (step 1Aiii). **c**, Cassettes are incubated in xylene on a hot plate with a magnetic stir bar (step 1B). **d**, Biopsies are individually incubated in the staining solution in a Roto-Therm rotator (step 2B). **e**, Biopsies are individually cleared by incubating in ECi on an orbital shaker (Step 2D). **! CAUTION** Experiments using human tissues must follow appropriate institutional and governmental regulations with respect to informed consent. The images in this figure are of de-identified tissues provided by an institutional tissue bank and are not considered human-subjects research.

Following tissue processing, samples are imaged with nondestructive 3D microscopy. Light-sheet microscopy is an ideal platform for high-throughput 3D pathology, enabling rapid and gentle imaging of cleared specimens. We initially developed Path3D for use with open-top light-sheet (OTLS) microscopy. In an OTLS architecture, illumination and collection optics are placed below a planar sample holder such that specimens are imaged from below, much like a flatbed document scanner. This approach is specifically optimized for pathology specimens of arbitrary size and shape^9–14^. Many other custom-built light-sheet systems exist ^15–18^, as do a number of commercial options (e.g., SmartSPIM, Leica SP8, Zeiss Lightsheet 7, Luxendo LCS SPIM/MuVi SPIM, Olympus/PhaseView Alpha3). Of particular note are single-objective light-sheet variants, which maintain a similar open-top geometry^19–29^. We use light-sheet microscopy in this protocol to illustrate key imaging considerations. However, our pipeline is in principle applicable to other 3D microscopy platforms (e.g., confocal microscopy, multi-photon microscopy). Further discussion of imaging platforms is provided in the ‘Comparison with other methods’ section. We provide strategies to establish the baseline performance of each microscope system for a given clearing protocol using 3D phantoms of diffraction-limited beads and an automated method to quantify a microscope’s point spread function (PSF, adapted from a method developed by Sofroniew [https://github.com/sofroniewn/psf], which is key for later quality control and troubleshooting (Fig. 4). We also offer guidelines for establishing ideal imaging conditions (considering detector counts, photobleaching, etc.).

**Fig. 4.**
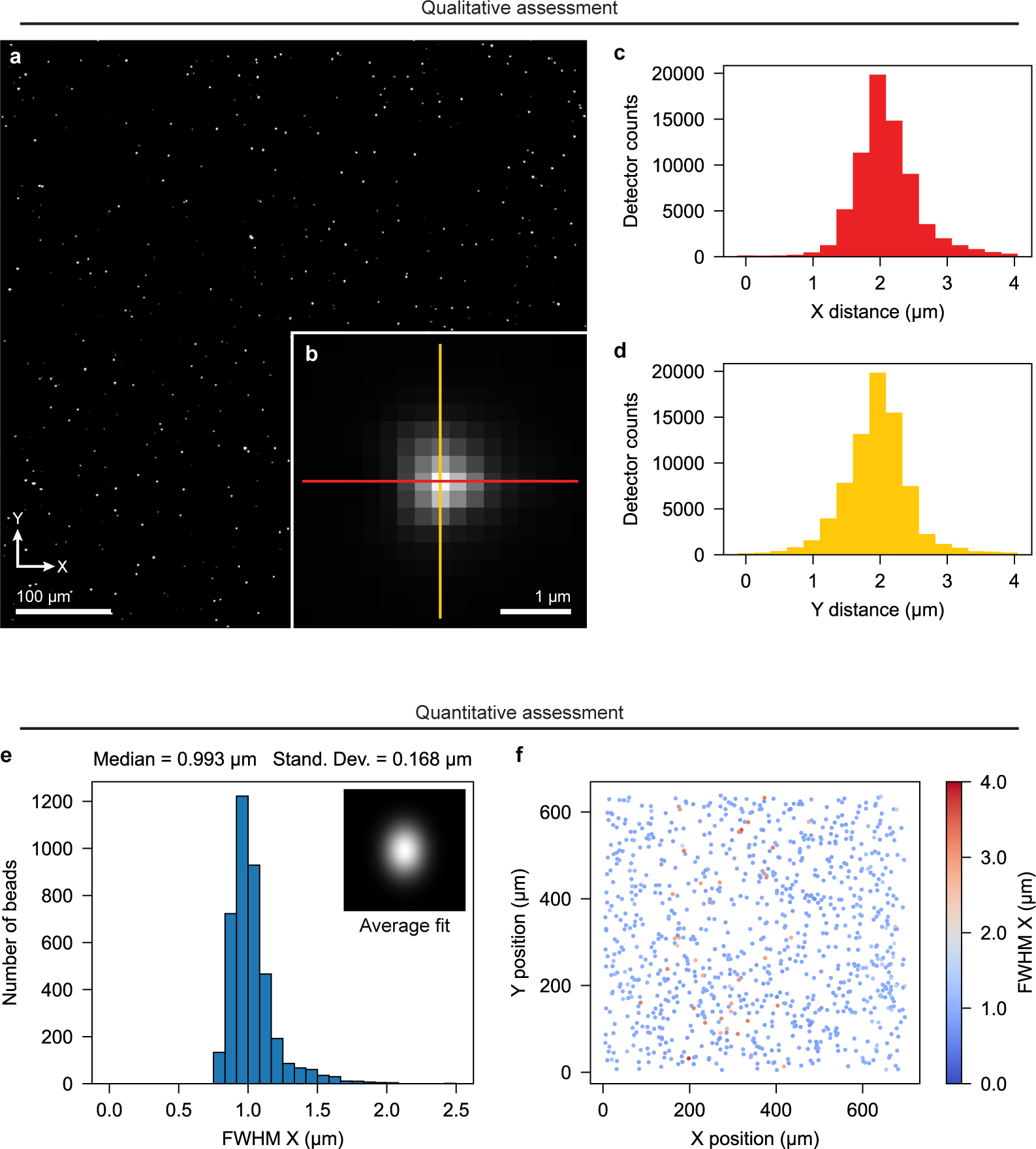
Example bead phantom and PSF data. Initially, agarose bead phantoms are qualitatively assessed in real time while being previewed to optimize the focus and alignment of the microscope (step 5). **a**, An example bead image (single slice) shows the relative bead density. Line profiles of a well-focused bead, **b**, in the X, **c**, and Y, **d**, directions show symmetric Gaussian profiles with peak detector counts well above the background level. After imaging a 3D volume, the Bead_PSF_computation package is used to quantify the full width at half maximum (FWHM) values for each bead (step 6). **e**, A histogram of lateral (X) FWHM values from all beads acquired with a well-aligned microscope) shows a distribution centered near the expected resolution with minimal spread. The inset shows an XY slice of the average 3D Gaussian curve fit. **f**, A heat map from a well-aligned example shows consistent lateral resolution with respect to X and Y position within the imaged volume. A perceptually uniform diverging colormap^93^ (Matplotlib^94^, “coolwarm”) is intentionally used to accentuate regions of particularly poor resolution. See Fig. S4 for additional details on heat maps.

For data processing and visualization, we focus on a fully open-source approach that can be performed by researchers with basic programming skills in Python. For nearly all 3D microscopy platforms, volumetric datasets of millimeter-to centimeter-scale tissues are first acquired by scanning the specimen in one primary direction (vertical or horizontal) to generate a 3D tile (a “volumetric brick”) of image data. Adjacent tiles are typically stitched and blended together to create a large contiguous volume before further image processing or visualization is performed. Various academic^30,31^ and commercial options (e.g., Aivia [https://www.aivia-software.com], Imaris [https://imaris.oxinst.com], Stitchy [https://www.translucencebio.com/products/stitchy]) exist for reconstructing 3D volumes from adjacent tiles. Likewise, a number of file formats have been developed for storing 3D datasets including HDF5^32^, Zarr^33^, OME-Zarr^34^, and N5^35^. This protocol utilizes an open-source method of aligning and reconstructing tiles (‘stitching and fusion’) using BigStitcher^36^ with HDF5^32^, which can be run as a FIJI^37^ plugin and requires minimal manual tuning of parameters.

Following reconstruction, datasets may be false-colored to mimic the pink and purple appearance of standard H&E histology. While anatomic pathologists are skilled at interpreting chromogenic (absorption-based) H&E slides, most are not experienced with interpreting multi-channel greyscale or arbitrarily colored fluorescence images. A variety of false-coloring methods have been developed, including machine-learning-based style-transfer methods^38–42^. While these strategies are appealing to convert images from one contrast mechanism to another, such as converting unstained whole slide images into H&E-like images^43^, they require large amounts of training data and are often hard to generalize to new data types (e.g. imaging conditions or tissue types not represented in the training data). Because Path3D datasets contain direct fluorescent analogs to conventional hematoxylin and eosin, we prefer a deterministic physics-based approach that models the absorption of light by conventional H&E dyes^44^. The explainability of this digital staining strategy is particularly appealing in the context of clinical studies and adoption by pathologists. Our approach is implemented as an open-source Python package, FalseColor-Python^45^.

Compared to viewing standard 2D whole-slide images, viewing and navigating 3D pathology datasets is not straightforward nor standardized. Many clinical or preclinical studies require interpretation of 3D datasets by panels of pathologists and researchers, so an easy-to-navigate visualization and annotation method is often desired. Here we offer guidance for using Napari, an open-source Python-based package for 3D image visualization (Fig. S9)^46^. In our experience, Napari provides relatively smooth and user-friendly visualization of 3D pathology data and offers built-in annotation features. Critically, Napari can dynamically load pyramidal file formats to minimize the amount of memory needed to navigate large 3D datasets. For users familiar with Python, Napari also offers additional analysis tools and freedom to write custom Python plugins. Other open-source visualization packages include QuPath^47^, which is designed for digital pathology and is ideal for viewing individual 2D slices in a 3D dataset, and FIJI (ImageJ)^37^, which is a widely used general-purpose tool for scientific image visualization. Commercial tools like Imaris [https://imaris.oxinst.com], Arivis [https://www.arivis.com], and Aivia [https://www.aivia-software.com] provide excellent volumetric (object rendering) as well as slice-by-slice visualization and annotation capabilities along with the professional technical support that many researchers value (Fig. 2a-b, Video S1). Recent academic studies have also explored novel methods for volumetric rendering and visualization of 3D pathology data^48,49^. For large studies with unique visualization requirements, custom interfaces (e.g., web-based) can also be developed.

Throughout the 3DPath protocol, continuous dataset evaluation and iterative optimization are essential to ensure that datasets are of consistently high quality for large-scale clinical or preclinical studies. Quality control is intentionally included as a routine step in the protocol rather than as a supplementary option, and we offer quantitative methods to assess resolution, contrast, and uniformity. Finally, we provide rough recommendations for the level of data quality that is necessary to produce high-quality false-colored imaging results that meet pathologists’ expectations. Our metrics also generally reflect the level of quality needed for effective downstream computational analyses.

### Applications and limitations

We have used Path3D to explore a number of clinical use cases for 3D pathology to date (Fig. 2). For example, various iterations of the protocol have been used to examine and quantify the 3D morphology of prostate cancer specimens^3,4,11–14,45,50^. This includes a proof-of-concept demonstration that 3D pathology can reduce certain ambiguities when performing Gleason grading of prostate cancer (which heavily influences treatment recommendations)^11^ and preliminary studies showing that the computational segmentation and analysis of prostate glands and nuclei in 3D have advantages for risk stratification^3,4^. Recently, we used a variation of Path3D to image whole lymph nodes from breast cancer patients and demonstrated that 2D pathology can underrepresent metastasis size compared to comprehensive 3D pathology^51^. We have also leveraged Path3D to show that diagnosis with deep-learning-assisted 3D pathology can detect esophageal neoplasia with higher per-biopsy sensitivity than standard 2D histology while also reducing pathologist workloads^5^. Additionally, the protocol has been used to generate 3D image atlases for various tissue types and disease states^52,53^. Larger studies comparing 2D and 3D pathology are underway, considering both inter- and intra-observer variability and overall accuracy (i.e., pathological grade vs. clinical outcome), and there are abundant opportunities to explore additional clinical applications of 3D pathology. Finally, the Path3D protocol has enabled a variety of preclinical studies such as characterizing 3D mouse liver cell cultures for microfluidic drug screening^54^, evaluating dissolvable microneedle arrays for transdermal drug delivery in mice, and imaging whole mouse kidneys at different scales to count and assess glomerular structure and function (Fig. S1c).

Manually reviewing 3D datasets using conventional workflows built around 2D slides can be labor-intensive, suggesting novel 3D-based review processes will be needed. Designing clinical studies that effectively use the large data volumes available from Path3D requires careful planning, and while not described here, computational analysis in conjunction with, or instead of, manual review is often appropriate. It is also critical to establish a robust data management plan, as raw datasets can be up to a terabyte (TB) in size (depending on image volume, resolution, etc.). Fortunately, there are many options for cloud-based storage and computational processing, which should be decreasing in cost over time. We have successfully used Path3D datasets for a number of computational use cases including image translation and glandular segmentation^3^, nuclear segmentation^4^, and deep learning-based triage^5^, and there are likewise opportunities to adapt computational techniques developed for 2D digital pathology to 3D datasets^55^.

Regardless of the specific staining and clearing protocol used (TO-PRO-3/eosin and ECi here), our data-processing and false-coloring workflow can be applied to generate H&E-like images from any fluorescent dataset with distinct nuclear and cytoplasmic channels. For example, we have used SYTOX-G or propidium iodide and NHS-ester staining with CUBIC-HV clearing to generate false-colored 3D datasets of human lymph nodes (Fig. S1a)^51^. We have also combined ECi clearing with fluorescent labeling of abundant reactive entities (FLARE) staining as an H&E analog^56^. Furthermore, we have shown that our workflow can be adapted for single-channel (acridine orange) datasets that are computationally separated into nuclear and cytoplasmic channels (Fig. S1b)^57^. Our focus here is on generating datasets that replicate standard H&E stains, the most commonly used by pathologists, meaning that Path3D datasets show tissue structure but do not on their own provide molecular information. However, we and others have shown that deep learning-based virtual-staining methods can be used to mimic certain targeted stains (e.g., immunolabeling) based on H&E-like datasets^3,39^.

It is important to note that while conventional H&E slides can be reimaged indefinitely, Path3D specimens, like any fluorescently labeled tissue, are subject to eventual photobleaching. The exogenous dyes used here are relatively bright (compared to endogenous fluorescent proteins, for instance) and photobleaching during initial imaging rounds is usually not an issue for well-stained specimens imaged at modest illumination powers (a few milliwatts of optical power measured at the sample). Nonetheless, repeated imaging of the same region will eventually lead to a significant drop in signal level. To avoid photobleaching valuable specimens with multiple redundant imaging rounds, it is important to ensure staining and imaging conditions are well-optimized before starting a large clinical study and that imaging regions are judiciously selected. While we have not attempted stripping and restaining Path3D specimens, previous reports suggest this may be possible^58,59^.

### Comparison with other methods

One alternative to non-destructive 3D microscopy of cleared tissues is automated serial-sectioning microscopy, in which thin tissue sections are rapidly cut from a thick specimen such that 2D images can be obtained throughout the specimen^60–64^. While serial sectioning obviates the need for chemical clearing, it is destructive of the tissue, which is not ideal for valuable clinical specimens, and it can introduce sectioning artifacts and complicate the reconstruction of 3D volumes. Non-destructive Path3D enables downstream analyses such as conventional 2D histology and molecular assays, which will be key to demonstrating the clinical value of 3D pathology and ultimately incorporating it into clinical workflows. Note that preliminary studies have shown that the H&E-analog and ECi-clearing steps described in this protocol do not adversely affect downstream standard pathology workflows (histology and molecular assays) and we are continuing to perform studies to rigorously demonstrate the “gentleness” of our methods when used on standard formalin-fixed specimens^5,52,65^.

There are several alternatives to the labeling protocols chosen for Path3D. Antibody stains are the primary alternative to small-molecule stains for clinical tissues, where genetically expressed endogenous labels are not possible. While antibodies are the gold-standard in terms of molecular specificity, the diffusion of large (∼150 kDa) antibodies into thick tissues can be extremely slow. Staining protocols that rely on passive antibody diffusion can take several weeks for tissues that are only hundreds of micrometers thick and tend to be complex compared to those using small molecule dyes^3,56^. A variety of methods have been developed combining immunolabeling and clearing, including strategies to speed penetration in thick tissues^66–71^. Variations of Path3D using antibodies instead of (or in addition to) small molecule dyes are possible, and while many antibody clones are compatible with ECi- and iDISCO-based clearing methods [https://idisco.info/validated-antibodies], combining immunolabeling and clearing can require specifically tailored protocols^8,72–74^.

It is also worth noting that a wide range of clearing protocols have been developed that vary in complexity, protocol duration, fluorophore compatibility, and achievable level of transparency^75,76^. Aqueous clearing methods are the primary alternative to solvent-based clearing approaches. These protocols tend to be gentler on the tissue and compatible with a wider range of fluorescent labels (especially endogenous markers). However, aqueous methods come with some practical challenges especially during extended imaging sessions. These include the tendency to form precipitates, which can be problematic for immersion-based microscopes, and the gradual evaporation of the water in the tissue and/or immersion bath, which can deteriorate tissue clarity and lead to refractive index mismatch issues. We have found that ECi-based clearing (used in our Path3D protocol) is a simple, fast, and reliable method for most tissues for imaging depths of up to a few millimeters. For applications where ECi clearing is inadequate, aqueous protocols like CUBIC^77–79^ can work well, as can more-elaborate solvent-based approaches, some of which involve detergents or delipidation steps^74,80^. An important note is that ECi-based clearing results in some degree of tissue shrinkage^6^ due to ethanol-based dehydration, which can have both advantages (modest reduction in imaging time) and disadvantages (modest reduction in effective spatial resolution).

Finally, there are several alternatives to light-sheet microscopy for nondestructive imaging of thick tissues. Initial studies used point-scanned techniques such as confocal and multi-photon microscopy^81–86^. Commercial point-scanned systems are widely available in core facilities, where dedicated staff and established imaging protocols can be leveraged to quickly establish feasibility in proof-of-concept studies. A major drawback to these laser-scanned systems is relatively slow imaging speed. In addition, some laser-scanned modalities, especially confocal microscopy, are highly light-inefficient, as they only collect a small fraction of the total excited fluorescence. In the most-basic form of light-sheet microscopy, a 2D plane within the specimen is selectively illuminated and imaged with a high-speed scientific CMOS camera, such that there is very little wasted light and, consequently, reduced photobleaching of the sample. By acquiring datasets plane-by-plane (as opposed to point-by-point or line-by-line), light-sheet systems can also achieve significantly higher imaging rates than laser-scanned microscopy systems and are ideal for high-throughput imaging of large tissue specimens. Depending on the system, light-sheet microscopes can, however, have more complicated sample-mounting workflows than commercial point-scanned systems.

### Experimental design

Here we outline several considerations when designing studies using our Path3D protocol. We specifically focus on large studies where tens to hundreds of samples must be processed and imaged, and where large data volumes must be managed.

#### Compatible tissues and selection of sample number and size

We have developed Path3D to work with formalin-fixed (FF) tissues as well as formalin-fixed paraffin-embedded (FFPE) tissues, which are first deparaffinized in the protocol. The procedure has shown success for a variety of clinical tissues including prostate^3,11–14,45,50,52^, esophagus^5,53^, lung^45^, kidney^45^, bladder, breast, pancreas, and colon (Fig. 2). We have also had success with preclinical specimens (mouse skin and kidney). For certain tissue types, we have found improved results with alternative staining and/or clearing protocols. As an example, lipid-rich lymph nodes benefit from CUBIC-HV clearing with aggressive delipidation to improve dye penetration (Fig. S1a)^51^. Additionally, i*n vivo* delivery of antibodies via retro-orbital injection allows for efficient and complete labeling of mouse kidney vasculature and glomeruli within minutes, even in the deepest layers of the organ (Fig. S1c). Nonetheless, Path3D works well for a wide variety of tissues and is the suggested starting point for new tissue types.

It is important to consider sample number and size when designing a study using 3D pathology. Tissue incubation times generally scale with sample size (the smallest dimension) and are typically longer than for 2D specimens to allow for diffusion of reagents into thick tissues. However, by obviating the need for mounting and sectioning by a skilled histotechnologist, the number of hands-on human hours is often less than or comparable to what is required for 2D histopathology. One should consider the tradeoff between tissue-preparation times and the amount of contiguous tissue volume that is needed for a selected analysis. For instance, the staining/clearing time for five 100-μm-thick specimens will likely be shorter than for a single 500-μm-thick specimen.

One should also note that imaging times and dataset sizes scale roughly linearly with tissue volume and to the third power with resolution. In other words, a 200 × 200 × 200 μm sample will, in general, take 8× longer to image and produce 8× more data than a 100 × 100 × 100 μm sample imaged on the same microscope. Likewise, imaging a sample with 1-μm resolution will take 8× longer and produce 8× more data compared to imaging with 2-μm resolution (given Nyquist sampling). Given that datasets can easily reach hundreds of gigabytes (GBs) and imaging sessions can take many hours for large samples, one should carefully consider what resolution and tissue volume are needed for a given study. In some cases, it may be preferable to image a smaller ROI rather than an entire specimen to reduce the total imaging volume. However, it is important to select ROIs with enough coverage to avoid reimaging (for instance if structures of interest are inadvertently truncated or missed) due to both the time required for reimaging and photobleaching concerns addressed above.

#### Computational and data storage requirements

The data processing workflow here was designed to run on a standalone workstation or local server running a standard Windows operating system set up with Python (e.g., Anaconda) but could in principle be adapted for cloud-based storage and processing. Fusing is the most computationally intensive step and benefits from a powerful CPU and large amount of RAM (for reference, we use a local server with 24 CPU cores (48 threads) and 662 GB of RAM). Stitching is much less computationally intensive but for convenience is often done with fusing on the same machine. Our stitching and fusing methods currently do not require a GPU. False coloring is less computationally intensive and can be run on a standard workstation with optional GPU acceleration. Napari performance is generally related to GPU specifications and RAM, though in most cases a standard workstation and GPU are adequate. For large studies, a large-capacity (∼100 TB) storage drive is necessary. Finally, compression schemes are worth considering. A variety of compression methods are available, including standard image compression algorithms (e.g., JPEG), popular lossless algorithms (e.g., BLOSC, ZSTD, GZIP), and certain lossy algorithms optimized for scientific applications (e.g., B3D^87^, SZ^88^). The ideal compression scheme for each study depends on factors such as the available storage and computational infrastructure, specific processing workflow, and microscopy platform.

Even with a powerful computer, processing times can be significant (often several weeks depending on dataset sizes and numbers). It is important to establish a streamlined workflow that requires minimal human input, such as batching datasets to run overnight. One should likewise consider how data will be transferred both internally (between local machines) and externally (to cloud-based resources, collaborators, etc.). Options include using a high-speed (10 Gbps) internal fiber network connection or loading data onto high-capacity portable drives that can be physically transferred or shipped. Regardless of strategy, transferring tens to hundreds of TBs of data is always a slow process, and we suggest planning workflows to minimize the number of transfers required.

#### Light-sheet microscopy platform

Factors to consider when selecting a specific light-sheet microscopy platform include volumetric imaging rate, ease of batch imaging (e.g., imaging successive samples without user intervention), maximum sample size, and maximum imaging depth (i.e., working distance). One should also consider the platform’s resolution, as this impacts both dataset size and imaging times as discussed previously. To use our stitching and fusing workflow, raw microscope datasets should be in a file format compatible with BigStitcher (e.g., HDF5). Alternatively, datasets can be stitched and fused using other methods (such as with native microscope pipelines) and converted to a standard format (e.g., HDF5, TIFF stack) for false coloring and/or visualization.

#### Visualization software

The most straightforward approach for image visualization by downstream users (e.g., pathologists), as described in this protocol, is to install display software (e.g., Napari, QuPath) and to load datasets onto a computer that can be accessed by users physically or via remote login. An alternative approach is to use a web-based viewer. While convenient, there are not yet open-source options for web-based viewing and this option is not discussed here. We do not recommend loading data and software directly onto every user’s computer given differences in hardware, installation requirements, etc.

### Required expertise

This protocol is intended for microscope end-users, including bioengineers, clinical researchers, and life scientists ranging from undergraduates to postdocs and staff scientists. As this is a multidisciplinary end-to-end workflow, no reader is expected to be an expert in all aspects of the process. For tissue preparation (steps 1 - 2), some experience with tissue staining and clearing is helpful but not required. For additional background, readers may consult previous work on clearing and staining, particularly ECi/iDISCO methods^6–8,75,89^. For sample imaging (steps 3 - 8), researchers should be competent using their chosen microscope platform (e.g., sample mounting, previewing samples, adjusting imaging parameters). For image processing (steps 9 - 20), users should have basic familiarity with Python (including package installation, development environments, Anaconda, etc.). Experience with using BigStitcher is also helpful; we recommend new users consult online tutorials [https://imagej.net/plugins/bigstitcher].

## Materials

### Biological materials

- Formalin-fixed (FF) or formalin-fixed paraffin-embedded (FFPE) tissue specimens **! CAUTION** Experiments using human or animal tissues must follow appropriate institutional and governmental regulations with respect to informed consent/care of animals. The images in Fig. S1b were obtained from excised specimens collected from patients undergoing head and neck cancer surgery at the University of Washington Medical Center. Approval was obtained from the University of Washington Institutional Review Board and patients provided informed consent. All other results from human tissues presented in this manuscript were from de-identified tissues provided by an institutional tissue bank and are not considered human-subjects research. The *in vivo* labeled mouse kidney tissues (Fig. S1c) were generously provided by Ruben M. Sandoval and Dr. Michelle M. Martinez-Irizarry at the Indiana University School of Medicine. These experiments followed National Institutes of Health Guidelines for the Care and Use of Laboratory Animals and were approved by the Animal Care and Use Committee at the Indiana University School of Medicine. The mouse flank skin tissues (Fig. S8b) were obtained from euthanized animals generously provided by veterinarians at an institutional animal facility at the University of Washington. These experiments therefore did not require approval from the University of Washington Institutional Animal Care and Use Committee.

### Reagents

- Ethanol (Decon Laboratories, cat. no. 2701) **! CAUTION** Ethanol is flammable and volatile, work with small amounts and store large quantities in a flammables cabinet. Ethanol is also an eye hazard. Wear appropriate PPE.
- Deionized (DI) water
- Sodium chloride (NaCl) (Spectrum Chemical, cat. no. SO160)
- 0.1N Hydrochloric acid (HCl) (Fisher Scientific, cat. no. SA54-1) **! CAUTION** Hydrochloric acid is a skin and eye hazard. Wear appropriate PPE.
- Xylene (Fisher Scientific, cat. no. X3P-1GAL) **! CAUTION** Xylene is flammable and volatile, work with small amounts and store large quantities in a flammables cabinet. Xylene is also a significant skin, eye, and respiratory hazard. Wear appropriate PPE and do not handle outside of a fume hood.
- TO-PRO-3 iodide, 1mM solution in DMSO (Thermo Fisher Scientific, cat. no. T3605) **CRITICAL STEP** TO-PRO-3 should be protected from ambient light during storage to avoid photobleaching.
- Alcoholic eosin Y 515 (Leica Biosystems, cat. no. 3801615) **! CAUTION** Alcoholic eosin is flammable and volatile, work with small amounts. Alcoholic eosin is also an eye hazard. Wear appropriate PPE. **CRITICAL STEP** TO-PRO-3 should be protected from ambient light during storage to avoid photobleaching.
- Ethyl cinnamate (ECi) (Thermo Fisher Scientific, cat. no. A12906.36) **! CAUTION** Ethyl cinnamate degrades polystyrene and other plastics. Use glass or polypropylene (PP) containers. Using incompatible plastic containers can cause the container to rupture or contaminate the contents.
- Agarose powder (Benchmark Scientific, cat. no. A1700)
- 150 nm diameter gold nanoparticles, stabilized suspension in citrate buffer (Sigma-Aldrich, cat. no. 742058)

### Reagent setup

#### Path3D staining buffer

The Path3D staining buffer is a mixture of 70% v/v ethanol in DI water with 10 mM NaCl at pH 4. To prepare 100 mL of staining buffer solution, add 70 mL ethanol and 58.44 mg NaCl to 10 mL DI water in a 250 mL beaker. Mix the staining buffer on the orbital shaker until the NaCl is fully dissolved. Add 0.1N HCl to the solution using a transfer pipette until the solution reaches pH 4 (typically 6 – 7 drops), measured using the pH meter. Finally, add DI water until the total volume of the solution is 100 mL. The buffer solution may be stored for one month at 4 °C but should be brought to room temperature (RT, ∼21 °C) before use.

**! CAUTION** Ethanol is flammable and volatile, work with small amounts and store large quantities in a flammables cabinet. Ethanol is also an eye hazard. Wear appropriate PPE.

**! CAUTION** Hydrochloric acid is a skin and eye hazard. Wear appropriate PPE.

### Equipment

- 250 mL beaker (Kimble Kimax, cat. no. 14000-250)
- Scale (VWR, cat. no. VWR-164AC)
- Orbital shaker (Labnique, cat. no. MT-201-BD)
- Transfer pipets (Fisher Scientific, cat. no. 13-711-9AM)
- pH meter (Thermo Fisher Scientific, cat. no. 13-645-611)
- Magnetic hotplate stirrer (VWR, cat. no. 97042-606)
- 1” magnetic stir bar (VWR, cat. no. 58948-138)
- Immersion thermometer (VWR, cat. No. 89095-628)
- Razor blades (VWR, cat. No. 55411-055)
- 6-well biopsy cassettes (Simport Scientific, cat. no. M503-10)
- Standard pencil (Ticonderoga, #2/HB)
- Tweezers (POLYPLAS Hameln GmbH, cat. no. 1404B-SA)
- Standard aluminum foil (Miktomeh, 1000 SQ FT)
- Standard micropipettes (Rainin, cat. no. HU Start Kit PL-LTS 20,200,1000u)
- 2.0 mL Eppendorf safe-lock tubes (Eppendorf, cat. no. 022363352)
- Roto-Therm rotator (Southern Labware, cat. no. H2024)
- 15 mL polypropylene conical tubes (Cellstar, cat. no. 188271)
- 50 mL beaker (Kimble Kimax, cat. no. 14000-50)
- 50 mL polypropylene conical tubes (Cellstar, cat. no. 227270)
- Light-sheet microscopy platform (see Experimental design section for details)
- Computing and data storage setup (see Experimental design section for details)

### Procedure

#### Tissue preparation

**CRITICAL STEP** The following procedure is optimized for 1- to 3-mm diameter punch-biopsy or needle-biopsy specimens (∼5 – 50 mm^3^ biopsy volume after trimming excess wax). Incubation times and reagent volumes may need to be adjusted based on sample size.

**! CAUTION** Experiments using human or animal tissues must follow appropriate institutional and governmental regulations with respect to informed consent/care of animals. The images in Fig. S1b were obtained from excised specimens collected from patients undergoing head and neck cancer surgery at the University of Washington Medical Center. Approval was obtained from the University of Washington Institutional Review Board and patients provided informed consent. All other results from human tissues presented in this manuscript were from de-identified tissues provided by an institutional tissue bank and are not considered human-subjects research. The *in vivo* labeled mouse kidney tissues (Fig. S1c) were generously provided by Ruben M. Sandoval and Dr. Michelle M. Martinez-Irizarry at the Indiana University School of Medicine. These experiments followed National Institutes of Health Guidelines for the Care and Use of Laboratory Animals and were approved by the Animal Care and Use Committee at the Indiana University School of Medicine. The mouse flank skin tissues (Fig. S8b) were obtained from euthanized animals generously provided by veterinarians at an institutional animal facility at the University of Washington. These experiments therefore did not require approval from the University of Washington Institutional Animal Care and Use Committee.

1 OPTIONAL Deparaffinization of FFPE specimens *(51 hrs)*

(A) Xylene preparation
  (i) In a fume hood, fill a 250 mL beaker with 100 mL xylene and heat to 60 °C (measured using an immersion thermometer) on a hot plate.
  (ii) Trim any large pieces of excess wax from FFPE specimens using a razor blade. **CRITICAL STEP** Use caution if trimming small or thin specimens as they are prone to cracking. Aggressive trimming is not necessary, and excess wax that is challenging to trim can be removed through longer xylene incubation in the following steps.
  (iii) Load FFPE specimens into 6-well histology cassettes (1 specimen per well, i.e., up to 6 specimens per cassette). Label cassettes with pencil or a xylene-resistant ink (Fig. 3b). 100 mL of xylene is adequate to process 1 or 2 cassettes. For more than 2 cassettes, larger volumes and/or multiple beakers may be needed to ensure adequate circulation of xylene.
  (iv) Add cassettes to the beaker of preheated xylene, ensuring they are fully immersed. Add a magnetic stir bar (Fig. 3c). **! CAUTION** Xylene is flammable and volatile, work with small amounts and store large quantities in a flammables cabinet. Xylene is also a significant skin, eye, and respiratory hazard. Wear appropriate PPE and do not handle outside of a fume hood. **CRITICAL STEP** Cover beaker with aluminum foil to prevent evaporation. **CRITICAL STEP** We do not recommend using cassettes other than standard histology cassettes (a durable polymer), as many plastics (such as polystyrene) are not chemically resistant to xylene and may dissolve.
(B) Deparaffinization
  (i) Adjust hotplate to maintain a constant xylene temperature of 60 °C (measured using the immersion thermometer) and incubate with stirring (∼ 200 rpm) for 48 hours.
  (ii) OPTIONAL Exchange xylene with fresh xylene after 24 hours for tissues with excessive amounts of wax.
(C) Ethanol wash
  (i) Turn off heat and let xylene cool for 15 – 20 minutes. Once xylene and cassettes have cooled to RT, remove xylene and add enough 100% ethanol to submerge cassettes (∼50 – 100 mL). Incubate with stirring (∼ 200 rpm) at RT for 1 hour.
  (ii) Exchange the ethanol with fresh ethanol and incubate for 1 more hour. **PAUSE POINT** Samples may be left in 100% ethanol for up to one week (in a sealed container to prevent evaporation).
  **TROUBLESHOOTING**
2 Staining with a fluorescent analog of hematoxylin & eosin (H&E)
  (A) Tissue rehydration *(3 hrs)*
    (i) In the 250 mL beaker, place cassettes in enough 70% ethanol (v/v solution in DI water) to submerge (∼50 – 100 mL) and incubate for a minimum of 3 hours on the orbital shaker with gentle agitation (∼60 – 80 rpm) at RT. **PAUSE POINT** Samples may be left in a sealed container submerged in 70% ethanol for up to one month.
  (B) Tissue staining *(2 days)*
    (i) Prepare staining solution of 1:500 v/v TO-PRO-3 and 1:100 v/v eosin (together) in Path3D staining buffer (see Reagent Setup).
    (ii) Transfer each specimen to a separate 2 mL Eppendorf tube and add 1.5 mL prepared staining solution to each tube. Incubate specimens for 48 hours in the Roto-Therm rotator with mixing mode at 20 rpm at RT (Fig. 3d). **! CAUTION** Alcoholic eosin is flammable and volatile, work with small amounts. Alcoholic eosin is also an eye hazard. Wear appropriate PPE. **CRITICAL STEP** Ensure that specimens are completely submerged in staining buffer during incubation. For this and all subsequent steps, fluorescently labeled specimens should be protected from ambient light (e.g., by covering tubes in aluminum foil) to prevent photobleaching. **TROUBLESHOOTING**
  (C) Dehydration *(2 hours)*
    (i) Transfer each specimen to a separate 15 mL polypropylene tube and add ∼10 mL 100% ethanol to each tube. Incubate specimens for 1 hour on the orbital shaker with gentle agitation (∼60 – 80 rpm) at RT.
    (ii) Exchange ethanol with fresh ethanol and incubate for at least 1 more hour.
    (iii) OPTIONAL Incubate large samples in 100% ethanol overnight to ensure complete dehydration. **CRITICAL STEP** It is important to remove all water in the specimen during this step as any residual water will impede clearing. Ensure that all sample holders and storage tubes are free from condensation and are completely dry, especially in humid environments. **CRITICAL STEP** Dehydrated specimens can be brittle. Be gentle if using tweezers to transfer specimens.
  (D) Clearing *(4 hours)*
    (i) Transfer each specimen to a new 15 mL polypropylene tube and add 10 mL ECi. Incubate specimens for 2 hours on the orbital shaker with gentle agitation (∼60 – 80 rpm) at RT (Fig. 3e).
    (ii) Exchange ECi with fresh ECi and incubate for 2 more hours. **! CAUTION** Ethyl cinnamate degrades polystyrene and other plastics. Use glass or polypropylene (PP) containers. Using incompatible plastic containers can cause the container to rupture or contaminate the contents. **PAUSE POINT** Cleared specimens may be stored in 100% ECi at RT for several weeks to months before imaging. Ensure that specimens are protected from light. We do not recommend storing cleared specimens in the refrigerator. **TROUBLESHOOTING**

### 3D microscopy

#### Characterize systems with optically cleared agarose bead phantoms

3 Make agarose nanoparticle phantoms *(1 hour)*
  (A) Fill the 50 mL beaker with 19 mL DI water. Add a magnetic stir bar and heat the liquid to ∼85 °C (measured using an immersion thermometer) on a hot plate.
  (B) Set stirring to a high speed (∼400 rpm) and add ∼1% wt/vol agarose powder (∼0.2 g for a 20 mL final volume). **CRITICAL STEP** Add the agarose powder slowly to prevent clumping.
  (C) Cover the beaker with foil and incubate with continued stirring until agarose powder is completely dissolved (∼10 – 15 min), adjusting the hotplate setting as needed to maintain a liquid temperature of ∼85 – 90 °C.
  (D) After dissolution, add ∼5% v/v 150 nm diameter gold nanoparticle suspension to the solution (∼1 mL nanoparticle suspension for a 20 mL final volume, yielding a target concentration of ∼2 × 10^5^ nanoparticles/mL). **CRITICAL STEP** Nanoparticles may settle to the bottom of the citrate buffer. Shake or vortex the container thoroughly before removing nanoparticles.
  (E) Turn off the heat and cool the mixture to 70 °C with continued stirring (10 – 15 min). **CRITICAL STEP** Keep the beaker covered with foil throughout agarose preparation and avoid overheating the mixture to prevent excessive evaporation. If too much water evaporates (more than ∼20%), the final agarose concentration will change and may make it hard to form or use phantoms. We do not advise preparing less than 20 mL agarose as a time as excessive evaporation becomes hard to avoid.
  (F) Pipette mixture into 2 mL microcentrifuge tubes (∼0.5 mL per tube). Close tubes and place lid-side down on a flat surface until cooled to RT (∼30 min). Remove each phantom and proceed with dehydration or transfer the phantoms to DI water for storage. Microcentrifuge tubes create ∼1 cm diameter cylindrical phantoms with flat bottoms, which are convenient for inverted or open-top microscopes with a planar sample holder (Fig. S2). However other mold shapes can be used to accommodate other sample mounting requirements. **CRITICAL STEP** Work quickly as the mixture will cool and solidify rapidly once it is in a small volume in the pipette or tubes. Close and invert each tube as it is filled to prevent the mixture from solidifying in the bottom of the tube. **PAUSE POINT** Uncleared phantoms may be stored indefinitely (several months or more) in DI water at RT. Many phantoms can be stored in the same container as long as they remain submerged. As nanoparticles are reflective as opposed to fluorescent, it is not necessary to protect phantoms from ambient light.
4 Dehydrate and clear agarose nanoparticle phantoms *(8 hours)* **CRITICAL STEP** For the following steps, multiple phantoms may be processed together. Use a minimum of ∼10 mL of liquid per phantom in each step (e.g., use 40 mL of liquid for 4 phantoms in a 50 mL conical tube), and ensure phantoms are fully submerged. The listed incubation times are minimums and longer incubations times will not adversely affect the phantom. Phantoms can be treated much like a fixed tissue and dehydrated/cleared in other media as needed.

(A) Incubate phantoms in 100% ethanol for 2 hours at RT.
(B) Exchange the ethanol with fresh ethanol and incubate for 2 more hours.
(C) Incubate phantoms in 100% ECi for 2 hours on the orbital shaker with gentle agitation (∼60 – 80 rpm) at RT.
(D) Exchange the ECi with fresh ECi and incubate for 2 more hours. **PAUSE POINT** Cleared phantoms may be stored indefinitely (several months or more) in ECi at RT. Many phantoms can be stored in the same container as long as they remain submerged.
5 Image agarose nanoparticle phantoms *(timing variable)* **CRITICAL STEP** Image phantoms in reflectance mode (i.e., no emission filter) with red light (e.g., 638 nm) illumination. The scattering signals from gold nanoparticles tend to be significantly brighter than fluorescent dyes, so it may be necessary to use a low laser power or place a neutral density filter in the illumination or collection light path to avoid saturating the detector.

(A) Preview the bead phantom and qualitatively assess the bead size and shape (Fig. 4a-d). Adjust the imaging parameters until the microscope is well aligned and beads are optimally resolved based on the following conditions:
  (i) Beads should have a signal to background ratio (SBR) of at least ∼20 (see Box 1). Peak signal counts should be well above the background level (i.e., signal counts in regions of agarose with no beads) and well below the detector’s saturation point. For a 16-bit detector with a background level of 100 - 300 detector counts, we aim for peak signal levels of at least 5,000 detector counts.

##### Box 1 Signal, background, and noise

Here, we define **peak signal level** as the average component of detector counts in brightly stained regions of the sample and **background level** as the average component of detector counts in unstained regions of the sample. The background level is a combination of detector background (e.g., dark counts), sample autofluorescence, and other sources of stray or ambient light. Background regions are typically straightforward to identify in the nuclear channel (any regions without nuclei). For the cytoplasm channel, identifying background regions can be challenging as eosin will stain most tissue components to some degree. For certain tissues, the gland or vessel lumens can be used to quantify the eosin background level. We define the **noise level** as the random variation (standard deviation) in the background of an image (Fig. S5).

We confirm adequate signal levels by checking the **peak signal level** and the **signal to background ratio (SBR)**, or “contrast” (which can be readily estimated by comparing the peak signal level to the background level), for each specimen. The random noise level tends to be dominated by detector noise in the context of Path3D (sCMOS-based light-sheet imaging) and is typically well below the target signal levels. Therefore, adequate peak signal level and SBR usually also implies adequate **signal to noise ratio (SNR)**, or “brightness”.
  (ii) For most systems, optimally resolved beads should form a symmetric (approximately Gaussian) point spread function (PSF) in all three dimensions. When previewing the phantom, view the beads in successive image planes (i.e., scrolling through a small volume) to qualitatively check the PSF shape and size in all three dimensions.
  (iii) For Nyquist-sampled imaging, the full width at half maximum (FWHM) of each bead should be ∼ 2 – 3 pixels in width. For some system geometries, the imaging plane seen while previewing may not be the highest resolution plane. Likewise in systems with anisotropic resolution, Nyquist sampling may not be expected in every spatial dimension.
  (iv) Bead size and shape should be consistent across the field of view in terms of region-to-region trends. Outliers and some variation between individual beads are expected (Fig. S3). **CRITICAL STEP** To accurately quantify resolution, beads should be diffraction-limited (i.e., 5× – 10× smaller than the microscope’s resolution). Smaller beads may be necessary for some imaging systems. However, if the goal is to ensure uniformity and not to accurately quantify the PSF, slightly larger beads are acceptable.
(B) Once beads are well-focused, collect a 3D volume (∼ 1 × 1 × 1 mm^3^). Select a volume that does not include the edges of the phantom or defects (cracks, air bubbles, etc.), as these can interfere with PSF quantification.
6 Quantify PSF dimensions across a 3D volume *(timing variable, computational time on the order of several hours)*

(A) Compute PSF dimensions using the Bead_PSF_computation python package (see Supplementary note 1 for additional details). **CRITICAL STEP** The provided package requires an input bead phantom dataset as a TIFF stack. In the computed results, the X and Y dimensions are in the TIFF image plane, while the Z dimension is orthogonal to the image plane.

(i) Install the Bead_PSF_computation package by running the following command from a git command line: git clone https://github.com/kevinwbishop/Bead_PSF_computation
(ii) Fuse the HDF5 dataset to a TIFF stack in BigStitcher, reorienting as needed (stitching is not necessary to quantify PSFs). See steps 9 - 12 for details.
(iii) Open `PSF_notebook.ipynb` in Jupyter and run on the fused TIFF stack. Step-by-step instructions are provided in the notebook file. **PAUSE POINT** Initially computing the PSFs is computationally intensive and can take several hours on a standard workstation. Once initial results are computed, the data can be saved by the notebook as a CSV file and loaded again later to quickly produce additional visualizations.
(iv) As a starting point, refer to the histograms (Fig. 4e) showing the X, Y, and Z dimensions (FWHM) of each bead’s PSF. For a well-aligned microscope, the median FWHM should roughly match the system’s resolution (within ∼25%), and the histogram should show an approximately normal distribution with minimal variance (< 25% standard deviation) in bead dimensions. The scatter plots (Fig. S5) and heat maps (Fig. 4f, S4) produced by the Jupyter notebook can subsequently be used to understand how the system’s resolution varies across the field of view in one or two dimensions. A well-aligned system should have global variations in FWHM of < 25% across the field of view (though bead-to-bead variance may be higher and local outliers will exist, Fig. S3). **TROUBLESHOOTING**
(B) Agarose phantom results provide a benchmark of the resolution achievable with the imaging system’s current alignment and are valuable for downstream troubleshooting. If the resolution measured with agarose phantoms is not within ∼25% of the microscope’s specifications, there is likely an alignment issue that needs to be addressed before proceeding.

#### Image tissue

*(Timing variable)*

**CRITICAL STEP** Due to variations in sample preparation, tissue structure, etc., some tuning of initial imaging parameters is typically required for each sample type to optimize image quality.

7 Mount the stained and cleared tissue on the microscope and preview the cytoplasm (eosin: 525 nm excitation peak, 545 nm emission peak) and nuclear (TO-PRO-3: 640 nm excitation peak, 655 nm emission peak) channels.
8 Adjust the microscope’s exposure time and laser power until signal levels are adequate, then image the tissue. Use the following steps to assess signal levels while previewing the specimen:

(A) Estimate the peak signal level and signal to background ratio (SBR) of each channel (see Box 1).
(B) For optimal false-coloring results, the SBR should be ∼10 or more. The peak signal level should be well above the image background level but below the detector’s saturation point in order to maximize dynamic range. For a 16-bit detector with background detector counts of 100 – 300, we ideally aim for peak signal levels of at least 2000 detector counts (SBR ∼10). Note that the required peak signal level and SBR will depend on the application. **CRITICAL STEP** Specimens are particularly prone to photobleaching while previewing the sample, where one may be continuously viewing the same plane in a specimen. We suggest starting with minimal laser power to initially identify a region of interest.

### Data processing

**CRITICAL STEP** In this section, we describe our overall data-processing workflow and offer practical guidance relevant to Path3D datasets. The exact programmatic steps will vary with each experimental and data-management setup. For detailed usage instructions and examples, users should consult the online documentation for each software package:

BigStitcher: https://imagej.net/plugins/bigstitcher

FalseColor-Python: https://github.com/serrob23/falsecolor

Napari: https://napari.org/stable

#### Image tile stitching and fusing using BigStitcher

*(Timing variable, computational time on the order of several hours)*

9 Install FIJI^37^ and the BigStitcher plugin^36^.
10 Load the raw dataset and inspect the image data under different views, regions, and magnification scales. Ensure that the image tiles are roughly aligned to one another and that image quality is relatively consistent across the dataset. **CRITICAL STEP** BigStitcher supports a variety of import file formats. Multiresolution pyramidal formats such as HDF5^32^ offer efficient navigation of large datasets, which is convenient for interactive previewing.
11 Run stitching (via the Stitching Wizard) to align the tiles based on image features, which can compensate for minor misalignments between raw image tiles. **TROUBLESHOOTING**
12 Run fusion to blend individual tiles into a continuous and seamless 3D volume. **CRITICAL STEP** The image fusion process can be very slow (several hours for 100 GB-scale datasets on a workstation with ∼512 GB of RAM). We recommend fusing the dataset on a workstation with a large amount of RAM (> 256 GB) for faster processing. Processing time and fused dataset size can be reduced by cropping the dataset to a subregion using the “Bounding Box” feature or reducing the resolution via downsampling.

(A) OPTIONAL When fusing to a TIFF stack file format, the slicing plane must be specified. Before fusing, use the “Interactively Reorient Sample” option to orient the sample view to lie in the desired slicing plane.

#### OPTIONAL Generating virtual H&E images using FalseColor-Python

*(Timing variable, computational time minimal)*

13 Install the Cuda Toolkit followed by FalseColor-Python. We recommend installing FalseColor-Python in its own virtual environment. **CRITICAL STEP** Many methods within FalseColor-Python (e.g., `fc.rapidFalseColor`) require a Cuda-enabled GPU. For use without a GPU, equivalent CPU-based methods (e.g., `fc.falseColor`) are provided.
14 All false coloring and image processing methods are contained in the coloring module of FalseColor-Python. Import this module using: `import falsecolor.coloring as fc` **CRITICAL STEP** Generating a virtual H&E image using FalseColor-Python requires two grayscale images: a nuclear channel (TO-PRO-3) and a cytoplasm channel (eosin), which will be referred to as “nuclei” and “cyto” below.
15 Read nuclei and cyto into memory using any standard image processing library such as Scikit-Image^90^, OpenCV [https://opencv.org], Tifffile^91^, or H5py [https://www.h5py.org]. Both the nuclei and cyto images should be NumPy arrays^92^ of the same lateral dimensions.
16 To improve the appearance of smaller structures within the final virtual H&E image, FalseColor-Python has an image-sharpening method, `fc.sharpenImage`. This method returns an image that has been convolved with a horizontal and vertical edge-sharpening kernel. For example: `sharp_nuclei = fc.sharpenImage(nuclei)` `sharp_cyto = fc.sharpenImage(cyto)`
17 Load nuclei and cyto color settings for the virtual H&E image using

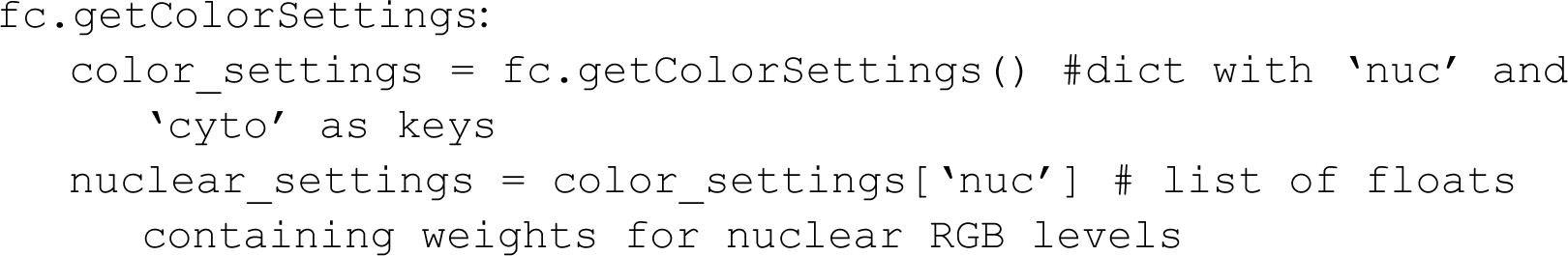
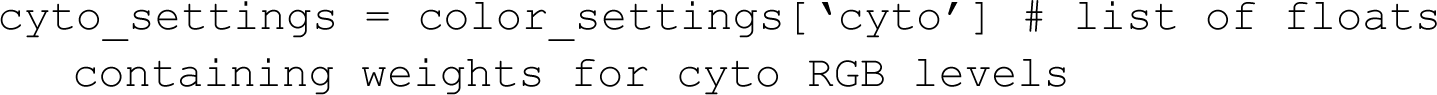 The color settings represent the weights that each channel (nuclei and cyto) contributes to the final RGB levels of the virtual H&E image. Default settings were empirically derived to match the hue and saturation levels seen in standard histology^45^. However, color preferences in histology are subjective and these weights can be adjusted to tune the coloration as desired. Each individual color level affects the relative color balance in the final H&E image. For example, nuclei with a bluer hue in the final virtual H&E image can be achieved by increasing the first constant in the `nuclear_settings` list. Further discussion of color settings and their effect on the final appearance of the virtual H&E image is available in the `Adjusting Color Settings.ipynb` notebook.
18 Generate a virtual H&E image from nuclei and cyto using `fc.rapidFalseColor: virtual_HE = fc.rapidFalseColor(nuclei, cyto, nuclear_settings, cyto_settings)` **CRITICAL STEP** `rapidFalseColor` has four additional input parameters: background levels for each channel (`nuc_bg_threshold` and `cyto_bg_threshold`) and normalization factors (`nuc_normfactor` and `cyto_normfactor`). These parameters are designated as integer values, which have been optimized for datasets in our lab but will likely change depending on the microscope and individual dataset. Each of these parameters affects the saturation levels of the final virtual H&E image in a slightly different way (Fig S7). The background threshold for each channel is the pixel value (assumed 16 bit) below which all pixels will be set to zero (and thus appear white in the final virtual H&E image). This can help reduce nonspecific background in empty regions of the image. Note that this level is subtracted from *all* pixels, and thus higher values for each channel will result in a lighter appearance in the virtual H&E image. The normalization factor for each channel is the constant that all pixels are divided by before the final coloration. This helps to reduce overly bright regions in the grayscale image such that they do not appear unrealistically saturated (or dark) in the final virtual H&E image. Further discussion of normalization settings and their effect on the final appearance of the virtual H&E image is available in the `Adjusting Color Settings.ipynb` notebook.

#### Visualizing fused and (optionally) false-colored datasets using Napari

*(Timing variable)*

**CRITICAL STEP** While Napari can be used out of the box on a standard workstation to view small datasets (i.e., smaller than the available RAM), viewing larger datasets can be inefficient or impossible. To view large datasets, users should use dynamic loading and/or compressed datasets to reduce computational requirements (Supplementary note 2).

19 Install Napari^46^ using Pip (we suggest the “batteries included” installation) in a new virtual environment.
20 Launch Napari and load the data Z-stack (raw fluorescence data or false-colored data). The Napari viewer allows users to pan, zoom in and out, and scroll in the Z-dimension through the dataset. Annotations can be added interactively using the “Layers” feature. **CRITICAL STEP** For visualization in Napari, false-colored datasets must be saved as an RGB TIFF stack. **CRITICAL STEP** Save annotations using “Save All Layers”. For 3-dimensional datasets, layers cannot be saved as a “napari SVG (*.svg)” file. Instead use “napari Save to Folder (*.*)”. The folder will include the original dataset and the coordinates of the shapes, points, and labels from each layer stored in comma-separated values (CSV) file. Load saved annotations by dragging and dropping the CSV file into Napari. **TROUBLESHOOTING**

### Quality control

*(Timing variable)*

**CRITICAL STEP** Ongoing quality control is essential to generating consistent data. The following steps should be used to iteratively establish the staining and imaging conditions at the start of a study, as well as to evaluate each collected dataset during a study. Quality control should be performed throughout an imaging study so that any problems can be caught and addressed promptly.

#### Quantitatively assess signal counts for each specimen

21 Fuse the raw HDF5 dataset as a TIFF stack sliced parallel to the tissue surface (HDF5 files must be viewed in BigStitcher, which does not allow for easy quantification of signal counts). See steps 9 - 12 for details. For large datasets, it is more efficient to fuse a few discrete Z levels of interest rather than the entire specimen. Accurate stitching is not necessary for initially assessing signal counts.
22 Plot line profiles across the specimen at multiple Z levels (e.g., near the tissue surface, 250 μm deep, 500 μm deep) (Fig 5). Based on qualitative assessment of images, select line profiles that adequately capture variations in signal counts (e.g., through a dimmer core, across intensity gradients from one side of the tissue to the other). When comparing across multiple specimens, select line profiles with similar positions and orientations as signal counts may be impacted by scan direction, light-sheet illumination direction, etc. Line profiles can be plotted in FIJI^37^ using the “Plot Profile” tool.

**Fig. 5.**
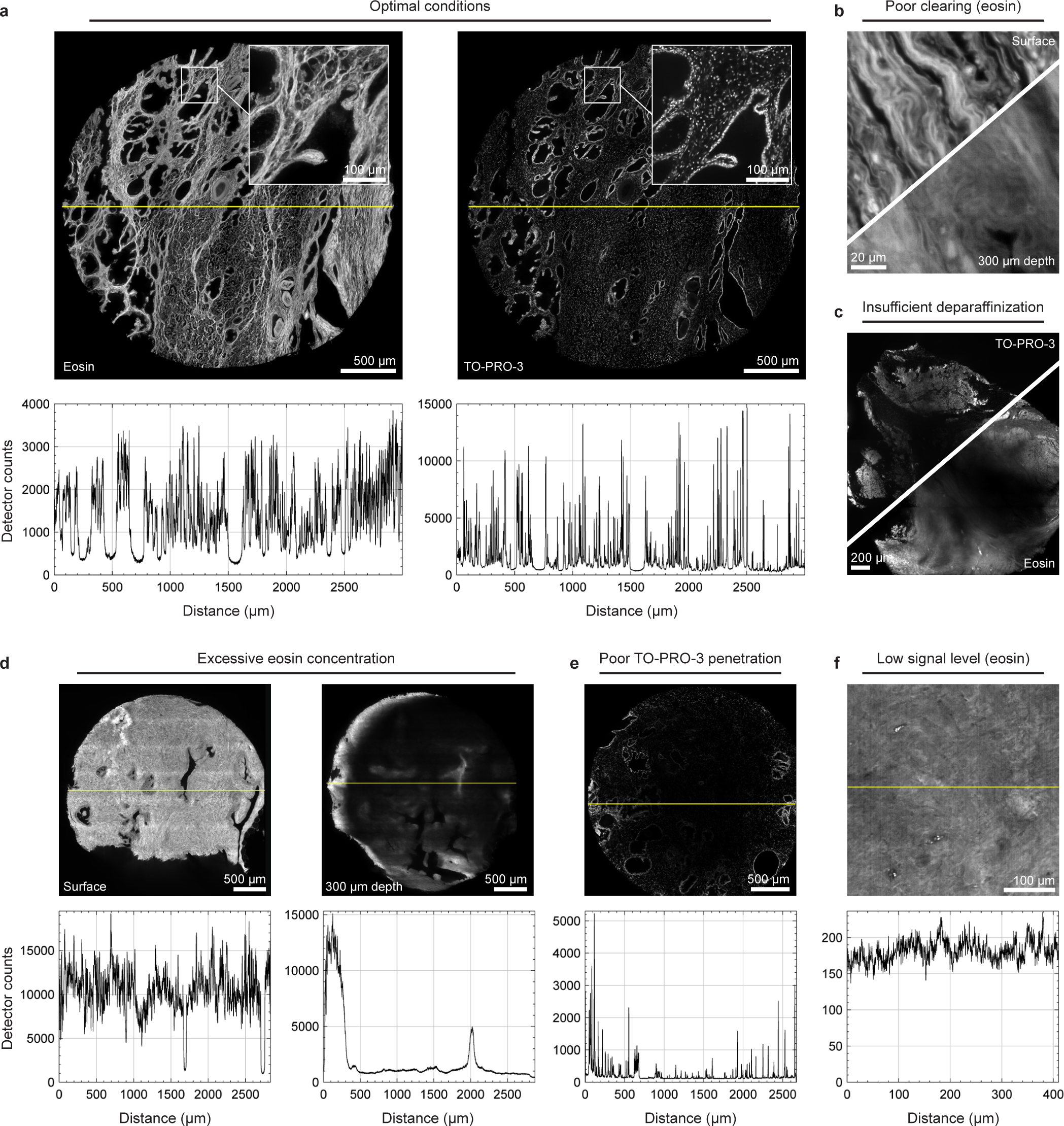
Examples of high- and low-quality tissue datasets. Images and line profiles (yellow line in each image) are from human prostate tissue imaged with OTLS microscopy and show various staining, clearing, and imaging conditions. **a**, Optimal staining and imaging conditions showing high SBR (> 10) and peak signal levels (> 2000 detector counts) in both channels with uniform signal levels across the tissue. Insets show well resolved collagen fibers (eosin channel) and nuclei (TO-PRO-3 channel). **b**, Poor clearing results in scattering of illumination and/or collection beams, degrading image sharpness with depth. **c**, Insufficient deparaffinization results in poor staining (especially at the core of the sample) in both channels. **d**, Excessive eosin concentration causes illumination light to be absorbed before reaching the interior of the tissue, resulting in bright edges and a dark core. Note the laterally asymmetric brightness when imaging at depth due to the illumination light entering the tissue from one side of the tissue in this particular imaging geometry (OTLS microscopy) (Fig. S7). **e**, Uneven staining due to poor TO-PRO-3 penetration in tissue resulting in bright edges and a dark core. **f**, Low eosin signal level (resulting in poor SNR) due to inadequate dye concentration, laser power, or exposure time. Note the presence of noise artifacts (vertical stripes) in the image due to insufficient SNR. **! CAUTION** Experiments using human tissues must follow appropriate institutional and governmental regulations with respect to informed consent. The results in this figure were from de-identified tissues provided by an institutional tissue bank and are not considered human-subjects research.
23 Confirm that the peak signal level and SBR are high enough without saturating the detector. See step 8 for additional details on assessing signal levels. **CRITICAL STEP** Datasets with inadequate signal levels can often be made to *appear* bright and well contrasted by tuning display settings in BigStitcher/FIJI. However, such datasets may not yield reliable false-colored results and may not be ideal for computational analysis. It is therefore essential to quantitatively assess signal levels using line profiles. **TROUBLESHOOTING**
24 Confirm that signals within each specimen are consistent both laterally and with depth. Ideally, peak signal counts for similar structures should vary by < 25% across a specimen (Fig. 5). **TROUBLESHOOTING**

#### Qualitatively assess image sharpness for each specimen

25 Assess image sharpness as a function of depth by qualitatively inspecting small features in the tissue architecture (such as isolated nuclei/subnuclear features in the nuclear channel or collagen fibers in the cytoplasmic channel). The degree of degradation with depth is driven primarily by the quality of optical clearing, and some degradation in sharpness is expected even for well-cleared tissues. For most tissue types, images should remain reasonably sharp at imaging depths of at least 500 μm. **TROUBLESHOOTING**

#### Approaches to troubleshooting image quality

**CRITICAL STEP** Loss of signal and loss of sharpness often occur together as they are related effects caused by refractive-index heterogeneities within the sample (leading to scattering and aberrations). However, in general, problems with staining or laser power/exposure time have a greater impact on signal counts, while problems with alignment and clearing have a greater impact on sharpness.

**CRITICAL STEP** For elusive image quality issues, we suggest sketching out the light paths of both the illumination beam and excited fluorescence for different regions of tissue. These details will vary based on the geometry of the specific imaging system, and understanding how light is traveling through the tissue to and from the imaging plane can provide insights into the source of a particular problem. For instance, low signal near a tissue edge where the illumination path length in tissue is significantly longer than the collection path length could be indicative of illumination attenuation rather than collection attenuation (Fig S8).

## Troubleshooting

Troubleshooting advice is available in Table 1.

**Table 1.**
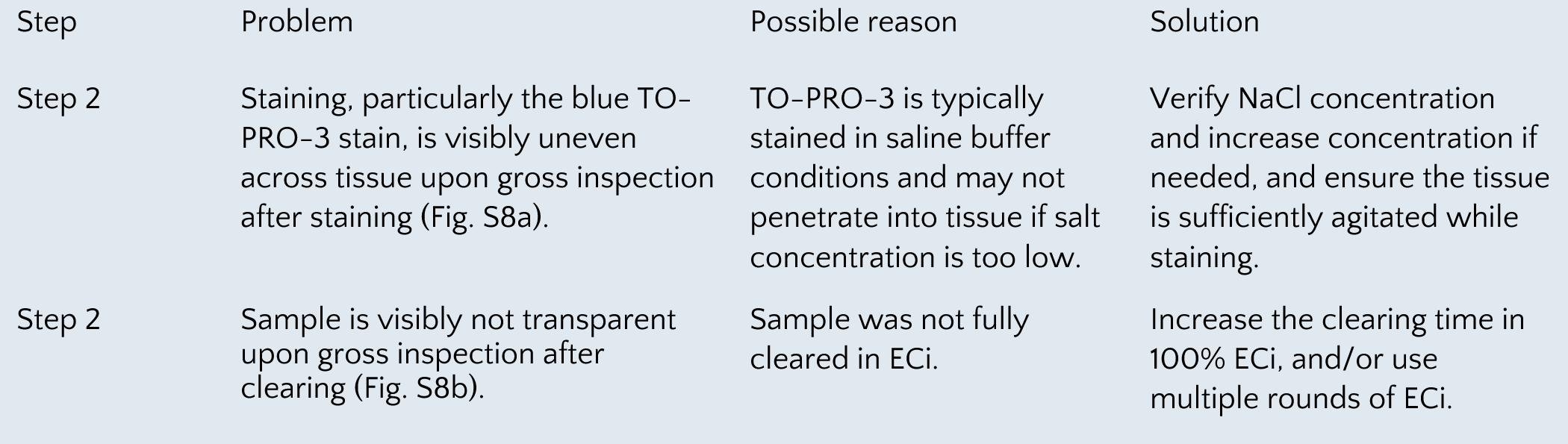
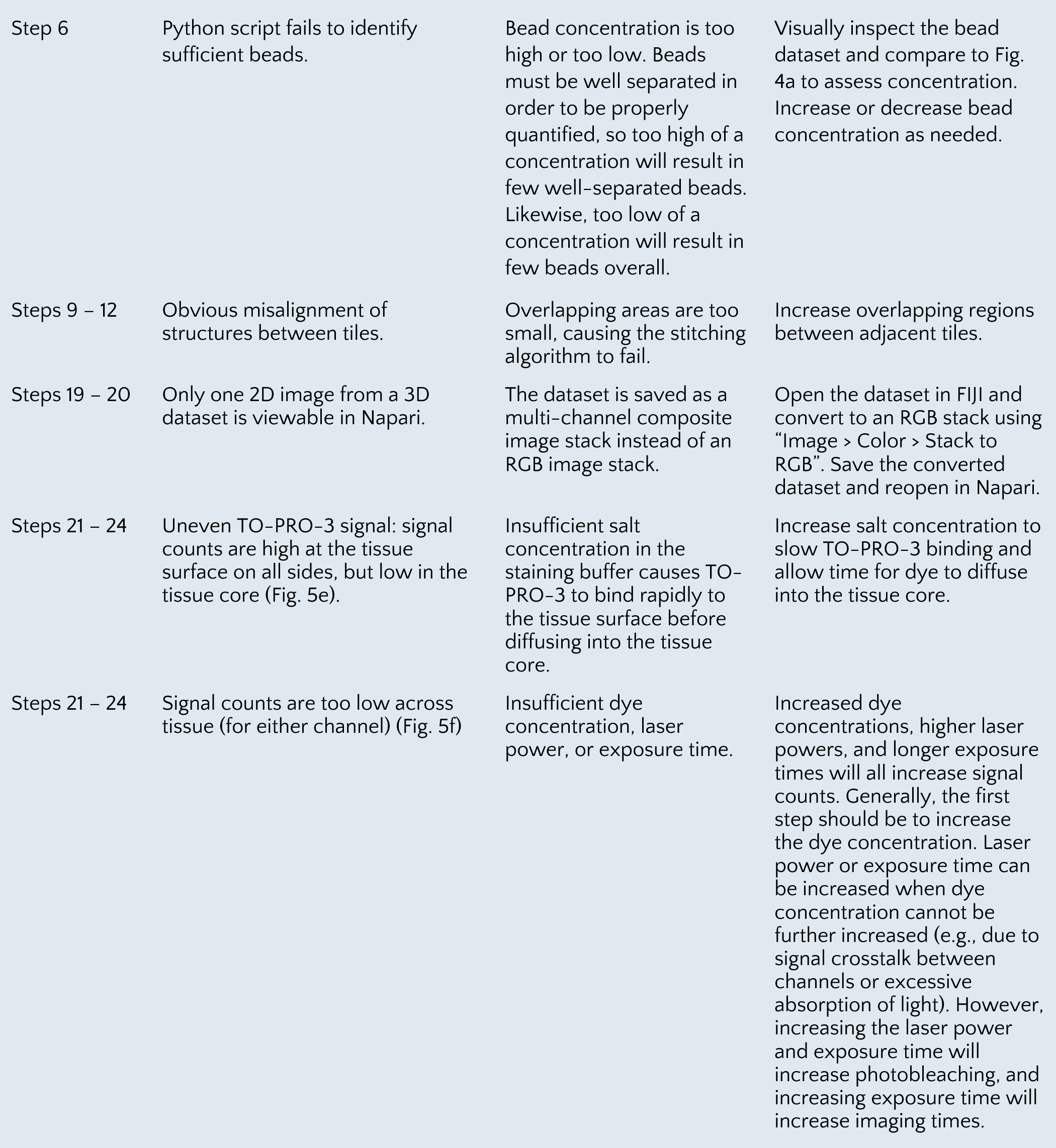
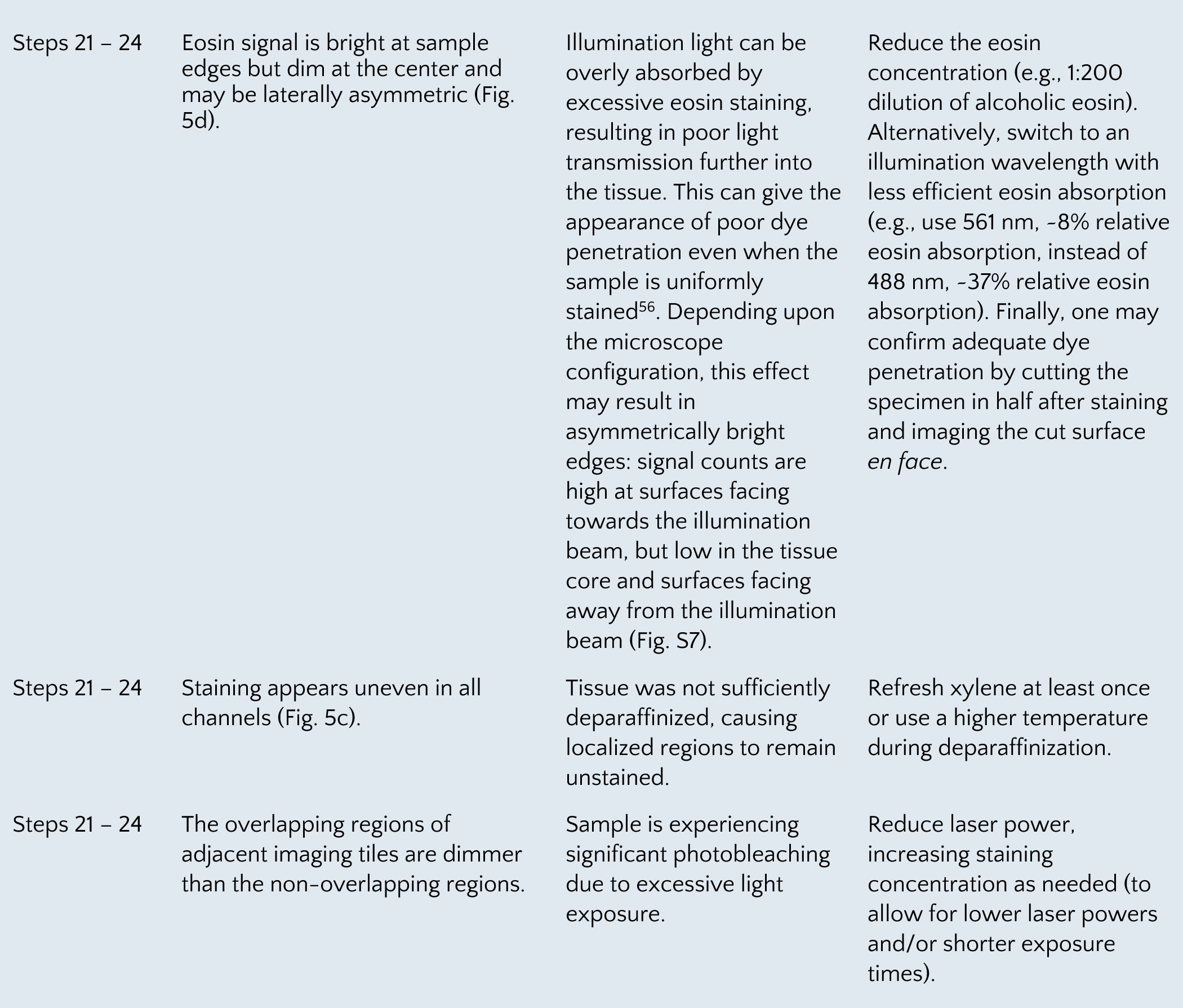
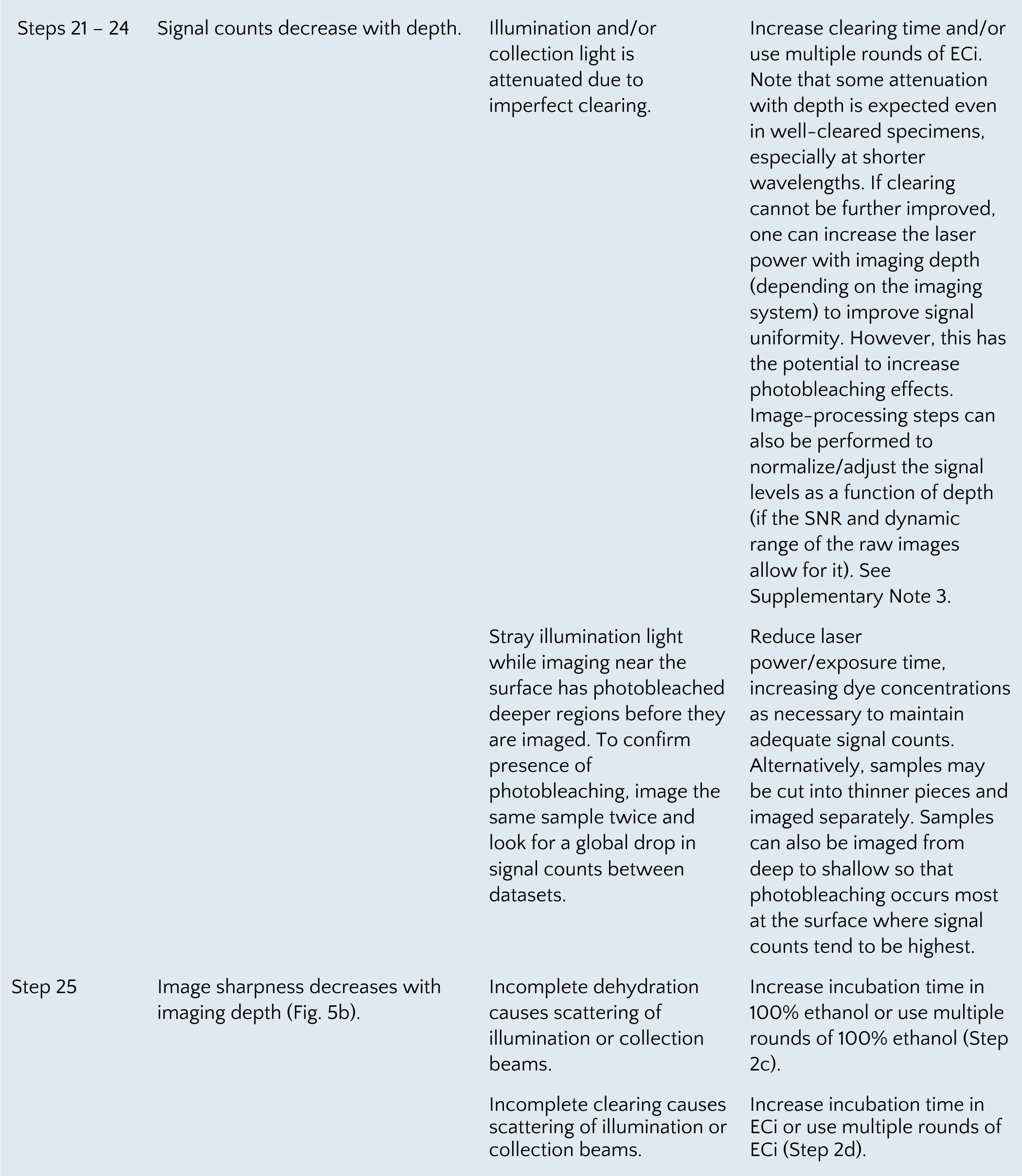
Troubleshooting of Path3D.

## Timing

Step 1 (optional), deparaffinization: 51 hours

Step 2, tissue staining and clearing: 57 hours

Step 3 - 4, agarose phantom preparation: 9 hours

Step 5 - 6, agarose phantom imaging and quantification: variable

Step 7 - 8, tissue imaging: variable

Step 9 - 12, stitching and fusing: variable (generally several hours)

Step 13 - 18, false coloring: variable (minimal computational time)

Step 19 - 20, visualizing data: variable

Step 21 - 25, quality control: variable

## Anticipated Results

We have employed Path3D on a range of tissue types including both human pathology tissues and preclinical animal tissues (both FF and FFPE). Here we present example results imaged with OTLS microscopy using both the standard Path3D protocol presented in this manuscript (Fig. 2) and protocol variants that have been adapted for specific use cases (Fig. S1). We also show examples of various visualization platforms that can be used to interact with Path3D datasets (Fig. S9 – S10).

## Archived human pathology specimens using standard Path3D

The standard Path3D protocol described here works well on a variety of archived human pathology specimens and is the suggested starting point for most tissue types. For example, we have used Path3D with prostate cancer specimens of various sizes^3,11–14,45,50,52^, including 3- mm punch (Fig. 2a) and 1-mm core-needle (Fig. 2b) biopsies. We have also used Path3D to image archived human squamous mucosa (Fig. 2c)^45^, colon cancer (Fig. 2d), and Barrett’s esophagus (Fig. 2e)^5^. Datasets are shown with both false coloring to mimic conventional H&E staining and with standard fluorescence coloring. Volumetric datasets allow both comprehensive assessment of tissue architecture (e.g. heterogeneous glandular morphology in prostate tissue, a key prognostic indicator for prostate cancer) and visualization of individual cellular and subcellular structures (e.g., subnuclear features) at high-resolution. Three-dimensional renderings (Fig. 2a-b, Video S1), orthogonal cross sections (Fig. 2b), or scrollable stacks of 2D images can be used to visualize three-dimensional tissue architecture.

### Path3D variants for clinical and preclinical tissues

For tissue types or use cases where the standard Path3D protocol is not ideal, variants using alternative clearing and/or staining methods can be employed. For example, human lymph nodes may be prepared with a protocol variant using a SYTOX-G or propidium iodide nuclear stain and an NHS-ester cytoplasmic stain with CUBIC-HV clearing (Fig. S1a)^51^. Volumetric renderings and cross sections taken at different depths show changes in tissue morphology with respect to depth that would not be apparent from a single 2D histology section. We also show an unfixed head and neck cancer human surgical specimen (tongue) imaged using a rapid single-stain variant (acridine orange) without clearing (Fig. S1b)^57^. In this case, a surface extraction algorithm was applied to generate a single 2D view of the irregular tissue surface from a thin 3D imaging volume. Nuclear and cytoplasmic channels were then computationally extracted from the single fluorescence channel and visualized with our standard false-coloring procedure. Finally, we show a whole mouse kidney with antibody labeling of vasculature and glomeruli using ECi clearing (Fig. S1c). Antibodies were delivered via *in vivo* retro-orbital injection to provide complete labeling within in a few minutes. This example uses a maximum intensity projection color-coded by depth to show kidney vasculature across the entire organ, an additional visualization option that works well for viewing sparse 3D structures as a single 2D image.

### Viewing and interacting with Path3D datasets

Path3D datasets can be generated in standard 3D image formats (e.g. TIFF, HDF5) for use with a range of existing 3D visualization tools. The Napari interface described above, being used to visualize a false-colored prostate specimen, is shown in Fig. S9. Figure S10a shows a false-colored esophagus specimen with overlayed pathologist annotations visualized using QuPath, an alternative open-source viewing method^5^. Custom visualization methods are also possible for specific applications. Fig. S10b shows a custom web-viewer developed by Ground Truth Labs being used to visualize a false-colored prostate specimen, which provides integrated grading tools to record specimen-level pathologist annotations.

## Data availability

Due to the large size of the imaging datasets shown in this manuscript, the datasets are not available in a public repository. They are available from the authors upon request.

## Code availability

All code used in this protocol is provided on GitHub. For license information, please see the corresponding GitHub page.

PSF computation: https://github.com/LiuBiophotonicsLab/Bead_PSF_computation

FalseColor-Python: https://github.com/serrob23/falsecolor

Normalizing signal levels: https://github.com/LiuBiophotonicsLab/FixImage3D

## Author contributions

L.A.E.B., Q.H., E.B., L.L., A.K.G., H.H., S.K., and J.T.C.L. developed the tissue preparation methods. K.W.B., L.A.E.B, A.K.G., and J.T.C.L. developed the 3D microscopy methods. Q.H., E.B., G.G., R.B.S., S.S.L.C., A.K.G., A.J., and J.T.C.L. developed the data processing methods. K.W.B., L.A.E.B., E.B., L.L., G.G., R.B.S., S.K., and J.T.C.L. developed the quality control methods. K.W.B., L.A.E.B, Q.H., E.B., L.L., C.P., G.G., R.B.S., A.K.G., H.H., and D.M. performed experiments and analyzed data. K.W.B., L.A.E.B., and J.T.C.L. led writing of the manuscript. All authors contributed to the manuscript.

## Acknowledgments

This work was supported by funding from the National Institutes of Health (NIH): R01EB031002 and R01CA268207 (J.T.C.L.); R00CA240681 (A.K.G.); and U01CA239055, 1R01LM013864, 1U01DK133090, U01CA248226, and 2R01DK118431 (A.J.); the National Science Foundation (NSF): 1934292 HDR: I-DIRSE-FW (J.T.C.L.) and NSF Graduate Research Fellowships DGE-1762114 (K.W.B. and L.A.E.B.); the Department of Defense (DoD) Prostate Cancer Research Program: W81XWH-20-1-0851 (J.T.C.L.) and W81XWH-18-10358 (J.T.C.L. and L.D.T); Prostate Cancer UK: MA-ETNA19-005 (J.T.C.L.), and the Pacific Northwest Prostate Cancer SPORE (P50CA097186).

The authors would also like to thank the following individuals for assistance with specimen collection and preparation for data presented in this manuscript: Suzanne M. Dintzis, Michael C. Haffner, Priti Lal, Chenyi Mao, Michelle M. Martinez Irizarry, Nick P. Reder, Deepti M. Reddi, Ruben M. Sandoval, Etsuo A. Susaki, and Inti Zlobec. Finally, the authors acknowledge the Canary Foundation for providing some specimens used in this study.

## Figure & supplementary files list

### Main text figures

1 Protocol overview

2 Image atlas showing Path3D datasets of archived human pathology specimens

3 Photos showing key tissue processing steps

4 Example bead phantom and PSF data

5 Examples of high- and low-quality tissue datasets

### Supplementary notes

Supplementary note 1: PSF computation details

Supplementary note 2: Improving Napari efficiency

Supplementary note 3: Image processing approaches to normalize signal level

### Supplementary figures

S1. Image atlas of clinical and preclinical tissues processed with variants of the standard Path3D protocol

S2. Photos showing agarose bead phantom preparation

S3. Example scatter plot illustrating local vs. global PSF variations

S4. PSF heatmaps of well-aligned vs. misaligned microscopes

S5. Identifying peak signal level, background level, and noise level

S6. False coloring settings

S7. Schematic of beam paths for troubleshooting

S8. Photos showing visual appearance of tissue after tissue preparation problems

S9. Napari interface for interacting with Path3D data

S10. Alternative interfaces for interacting with Path3D data

### Additional files

Supplementary video 1: Volumetric rendering showing a Path3D dataset of a 3-mm diameter punch biopsy of prostate cancer tissue false colored to mimic the appearance of standard H&E staining.

