## Supplementary information for "An end-to-end workflow for non-destructive 3D pathology"

[1] Department of Mechanical Engineering, University of Washington, Seattle, Washington, USA
[2] Department of Bioengineering, University of Washington, Seattle, Washington, USA
[3] Department of Biology, University of Washington, Seattle, Washington, USA
[4] Department of Chemistry, University of Washington, Seattle, Washington, USA
[5] Department of Biomedical Engineering, Emory University and Georgia Institute of Technology, Atlanta, GA, USA
[6] Department of Oncology, Division of Precision Oncology, University Hospital of Geneva, Geneva, Switzerland
[7] Department of Clinical Pathology, Division of Clinical Pathology, University Hospital of Geneva, Geneva, Switzerland
[8] Department of Laboratory Medicine and Pathology, University of Washington, Seattle, Washington, USA
[9] Department of Physiology and Biophysics, University of Washington, Seattle, Washington, USA
*

### Supplementary information contents

**Notes**

Supplementary note 1: PSF computation details
Supplementary note 2: Improving Napari efficiency
Supplementary note 3: Image processing approaches to normalize signal level

**Figures**

S1. Image atlas of clinical and preclinical tissues processed with variants of the standard Path3D protocol
S2. Photos showing agarose bead phantom preparation
S3. Example scatter plot illustrating local vs. global PSF variations
S4. PSF heatmaps of well-aligned vs. misaligned microscopes
S5. Identifying peak signal level, background level, and noise level
S6. False coloring settings
S7. Schematic of beam paths for troubleshooting
S8. Photos showing visual appearance of tissue after tissue preparation problems
S9. Napari interface for interacting with Path3D data
S10. Alternative interfaces for interacting with Path3D data

**Additional files**

Supplementary video 1: Volumetric rendering of 3 mm prostate punch

### Supplementary note 1 | PSF computation details

The Bead_PSF_computation package allows automatic computation of point spread function (PSF) dimensions across a 3D imaging volume. The input dataset is a 3D volume of diffraction-limited beads (e.g. gold nanoparticle embedded in agarose). First, beads centroids are located within the 3D volume using a method developed by Sofroniew [https://github.com/sofroniewn/psf]. Beads which are not well separated from one another, or whose PSFs are significantly truncated by the volume edges are excluded from further analysis. Next, a 3D Gaussian function is fit to each bead, which takes the following form:

$$G(x,y,z) = C + A * \mathrm{Exp}\left(-\left[\frac{(x-x_o)^2}{2\sigma_x^2} + \frac{(y-y_o)^2}{2\sigma_y^2} + \frac{(z-z_o)^2}{2\sigma_z^2}\right]\right)$$

where $C$ and $A$ are intensity offset and amplitude constants, respectively, $x_o$, $y_o$, and $z_o$ are expected values in each dimension, and $\sigma_x$, $\sigma_y$, and $\sigma_z$ are standard deviations in each dimension. Finally, the full width at half maximum (FWHM) is computed in each dimension for each bead's Gaussian fit. An arbitrary rotation can optionally be added if the expected PSF is rotated relative to the volume's coordinate system (note that this will impact the relative orientations of FWHM measurements).

Note that diffraction-limited beads are inherently barely resolved (i.e. 2 – 3 pixels per FWHM given Nyquist sampling) and significant variation in Gaussian fits and FWHM measurements between individual beads is expected. A properly aligned microscope should have a global variance (i.e. average FWHM values in one region relative to another) of 25% or less, while variance between individual beads may be significantly higher (Fig. S3).

The resulting data (i.e. centroid coordinates and FWHM values for every bead) can be plotted against one another using the package's three plotting functions:

Histograms (Fig. 4e) show the distribution of FWHM values in a given dimension without regard to the spatial position of each bead. Histograms are a useful starting point to assess the resolution variation within an acquired volume. Some outliers are expected, but most beads should be clustered around a central value.

Scatter plots (Fig. S3) show each bead as a single point and plot the bead's FWHM value in a given dimension relative to the bead's centroid coordinate in one dimension. Scatter plots are helpful to visualize changes in resolution as one moves across the field of view.

Heat maps (Figs. 4f and S4) show each bead as a single point, which is color-coded according to the bead's FWMH value in a given dimension and plotted based on the bead's centroid coordinate in two dimensions. Heatmaps are helpful for visualizing higher order changes in resolution. Beads are colored using the "coolwarm" colormap from Matplotlib[94], a perceptually uniform diverging colormap[93]. This choice is intentional in order to accentuate regions of particularly poor resolution.

### Supplementary note 2 | Improving Napari efficiency

Napari will by default attempt to load entire TIFF stacks into RAM for visualization, which can be inefficient for large datasets. Computational performance can be improved by instead dynamically loading subregions of a tiled pyramidal TIFF file as the user views them ("lazy loading"). This can be accomplished by using Tifffile[91] to read data as a Zarr format[33] into a Dask array [https://docs.dask.org/en/stable]. For example:

```
import dask.array
import napari
import zarr
import tiffile

store = tifffile.imread('dataset.tif', aszarr=True)
z = zarr.open(store, mode='r')
data = dask.array.from_zarr(z)
viewer = napari.view_image(data)
```

Loading time and storage requirements can also be improved by compressing TIFF files before storage. For example, Tifffile can be used to apply JPEG compression, where a 90% quality compression level can reduce file sizes by ~10× in some cases:

```
import tifffile

tf = tifffile.TiffFile('dataset.tif')

with tifffile.TiffWriter('compressed.tif', bigtiff=True) as tif:
    options = dict(metadata=None,
                   photometric='rgb',
                   tile=(512, 512),
                   compression=('jpeg',90))
    for page in tf.pages:
        tif.write(page.asarray(),**options)
```

### Supplementary note 3 | Image processing approaches to normalize signal level

Optimizing experimental conditions should be the first step in addressing nonuniform signal levels across a dataset. However, even after careful optimization, some variation in signal level across large datasets can be unavoidable. Image-processing steps may be applied to improve signal level uniformity if the SNR and dynamic range of the raw images allow. Here we describe computational approaches to two common problems, lateral variations across multi-tile datasets and drops in signal level with depth. The code snippets provided below are for illustrative purposes; complete implementations of these approaches are provided on GitHub:

https://github.com/LiuBiophotonicsLab/FixImage3D

**Lateral variations**

A variety of factors, such as photobleaching of the overlapping regions between adjacent tiles or illumination light intensity variations across the width of the light sheet can result in lateral variations in signal level across a dataset. These variations are often especially apparent at seams between tiles.

The `stripe_fix` function normalizes the lateral signal level across a dataset. First, the average horizontal line profile is computed across each Z level:

```
## Calculate profiles with background removed
line_prof_n_nobg = np.zeros((img_nobg.shape[1]), dtype = np.float64)

## Grab horizontal line profile
for col in range(img_nobg.shape[1]):
      line_prof_n_nobg[col] =
         np.sum(img_nobg[:, col].astype(np.float64))

## Normalize line profile
line_prof_n_nobg = line_prof_n_nobg / np.max(line_prof_n_nobg)
```

Then, each row of the Z level is divided by the line profile, leveling the intensity horizontally across the image. Any negative values (a result of background subtraction) are set to be zero:

```
for row in range(img_nobg.shape[0]):
        img_nobg[row, :] = img_nobg[row, :] / line_prof_n_nobg

## Set negative values to zero
img_nobg[img_nobg < 0] = 0
return img_nobg.astype(np.uint16)
```

**Depth variations**

Even in well-cleared specimens, light scattering or absorption can lead to some attenuation of light intensity as the imaging depth is increased. This results in a drop in signal levels at deeper layers relative to the tissue surface.

The `Fix_script.py` script corrects lateral and depth variations throughout the depth of a dataset. First, the 2nd percentile (`p2`) and 98th percentile (`p98`) of signal intensities are computed for every layer from an 8× downsampled dataset:

```
p2, p98 = np.percentile(img_8x,
                        (2, 98),
                        axis = (1,2))
```

Then, in each z-level, the lateral normalization function (`stripe_fix`, described above) and a contrast stretching function (`contrast_fix`) are applied. The contrast stretching function rescales the intensity profile of each layer to a consistent intensity scale, commensurate to the surface level, based on the calculated `p2` and `p98`:

```
for z in range(len(img_3d)):
    img_corrected[z] = stripe_fix(img[z])
    img_corrected[z] = contrast_fix(img_corrected[z],
                                    p2[z],
                                    p98[z])
```

The correction is performed independently on each layer, ensuring that the result of one layer remains unaffected by the others. This enables the possibility of parallelization to expedite the processing time.

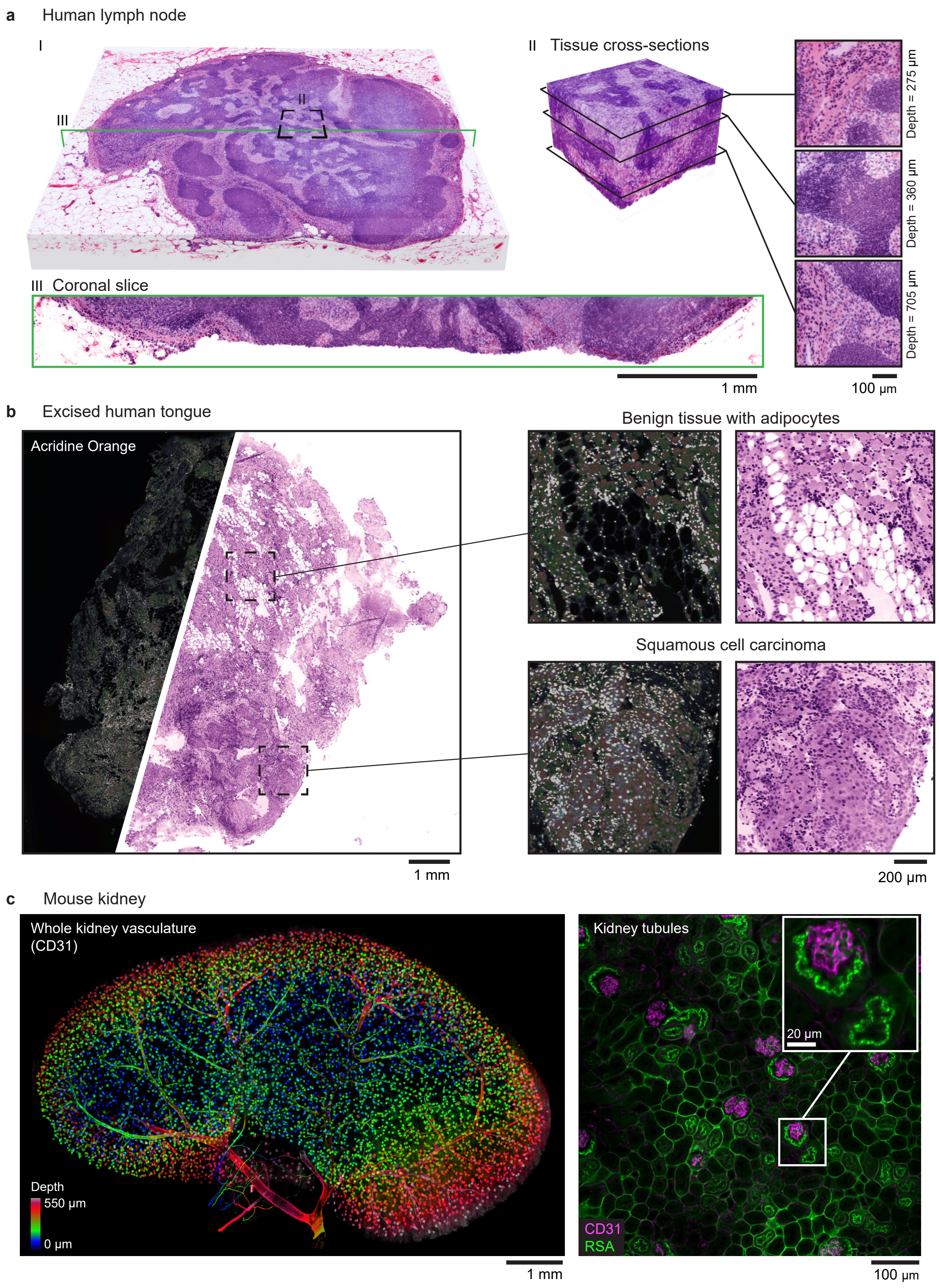

a
Human lymph node
I
II
III
II Tissue cross-sections
Depth = 275 µm
Depth = 360 µm
Depth = 705 µm
III Coronal slice
1 mm
100 µm
b
Excised human tongue
Acridine Orange
Benign tissue with adipocytes
Squamous cell carcinoma
1 mm
200 µm
c
Mouse kidney
Whole kidney vasculature
(CD31)
Depth
550 µm
0 µm
1 mm
Kidney tubules
20 µm
CD31
RSA
100 µm

**Fig. S1 | Image atlas of clinical and preclinical tissues processed with variants of the standard Path3D protocol.** **a**, Fixed human lymph node prepared with a CUBIC-clearing-based Path3D variant[51] and false colored to mimic the appearance of standard H&E staining, shown as volumetric renderings (generated using Aivia) and 2D cross sections. **b**, Fresh human head & neck cancer surgical specimen (tongue) prepared using a single-stain (acridine orange) Path3D variant without clearing. A single 2D view of the irregular tissue surface was computationally extracted from a thin 3D volume. Nuclear and cytoplasmic channels were computationally separated and false colored to mimic the appearance of standard H&E staining[57]. **c**, Whole mouse kidney with vasculature and glomeruli labeled with antibodies delivered via *in vivo* retro-orbital injection and cleared with ECi. A maximum intensity projection color-coded by depth (left) allows visualization of the whole kidney vasculature in a single image.

**! CAUTION** Experiments using human or animal tissues must follow appropriate institutional and governmental regulations with respect to informed consent/care of animals. The images in panel **a** were from de-identified tissues provided by an institutional tissue bank and are not considered human-subjects research. The images in panel **b** were obtained from excised specimens collected from patients undergoing head and neck cancer surgery at the University of Washington Medical Center. Approval was obtained from the University of Washington Institutional Review Board and patients provided informed consent. The tissues shown in panel **c** were generously provided by Ruben M. Sandoval and Dr. Michelle M. Martinez-Irizarry at the Indiana University School of Medicine. These experiments followed National Institutes of Health Guidelines for the Care and Use of Laboratory Animals and were approved by the Animal Care and Use Committee at the Indiana University School of Medicine.

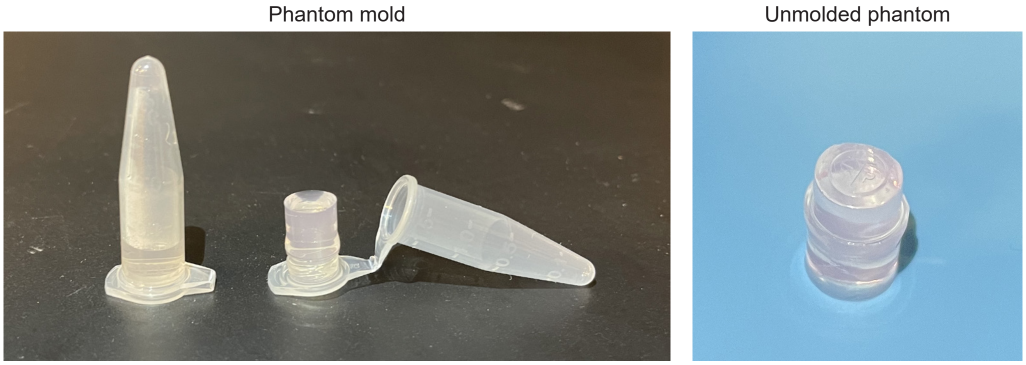


**Fig. S2 | Photos showing agarose bead phantom preparation**. Phantoms molded in inverted microcentrifuge tubes (step 3F) form flat-bottomed cylindrical plugs, which are convenient for use with inverted or open-top microscopes with a planar sample holder.

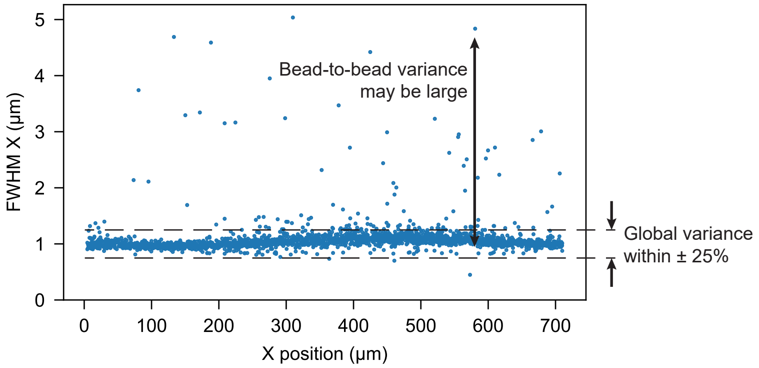


**Fig. S3 | Example scatter plot illustrating local vs. global PSF variations.** The plot was produced by the Bead_PSF_computation package (step 6) and shows individual beads in a volumetric agarose phantom dataset. Scatter plots show how a microscope's lateral resolution (FHWM of individual beads) changes across the field of view (X position). Well-aligned microscopes should have a global variance (i.e., average FWHM values in one region relative to another) of 25% or less. Due uncertainty inherent in measuring the FWHM of diffraction-limited beads, there may be significantly higher variance between individual beads.

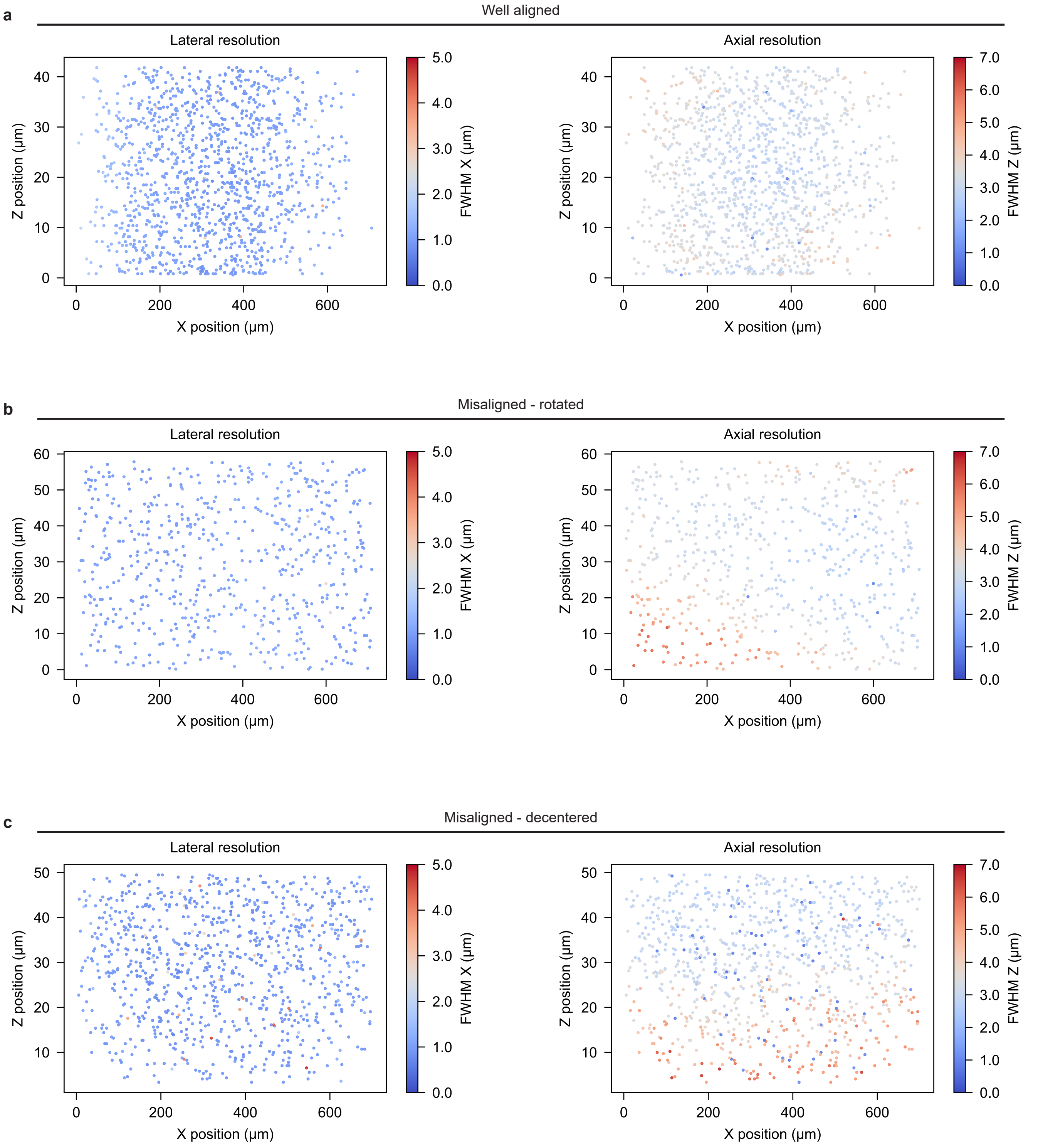


**Fig. S4 | PSF heatmaps of well-aligned vs. misaligned microscopes**. Heat maps were generated with the Bead_PSF_computation package (step 6) from example agarose bead datasets and show how a microscope's resolution varies across the field of view (FOV). Each point represents a single diffraction-limited bead. The bead's position indicates its location in the FOV (XZ plane in this case), while the color indicates the full width at half maximum (FWHM) of the fit Gaussian function in the lateral or axial direction. In the well-aligned case, **a**, there is minimal variation in resolution across the FOV. In the rotated case, **b**, the axial resolution is degraded in the lower left and upper right corners of the FOV, suggesting the light-sheet focus is rotated relative to the camera. In the decentered case, **c**, the axial resolution at the bottom of the FOV is degraded relative to the top of the FOV, indicating the light-sheet focus is not centered within the FOV. Note that a perceptually uniform diverging colormap[93] (Matplotlib[94], "coolwarm") is used, such that beads near the center of the FWHM range appear dull and beads at the edges of the range appear brightly colored. This was selected intentionally to accentuate regions of particularly poor resolution, as these regions are typically the most informative for assessing resolution uniformity.

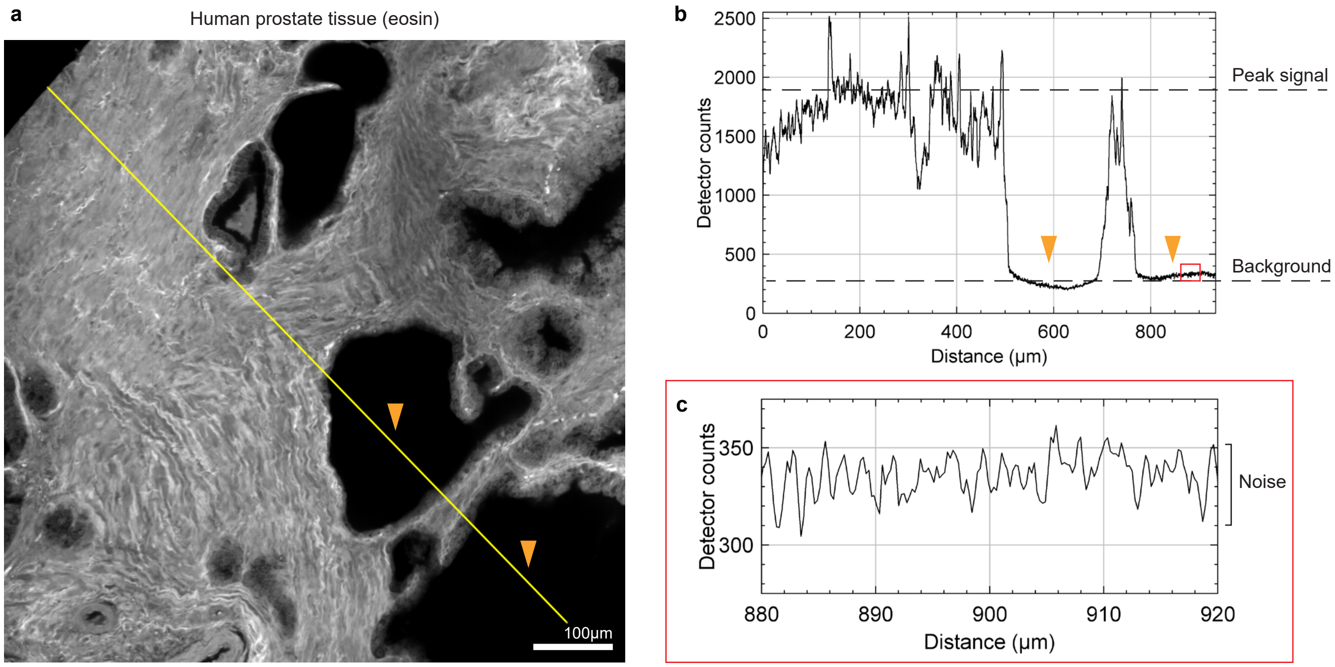


**Fig. S5 | Identifying peak signal level, background level, and noise level. a**, Human prostate tissue with eosin staining. **b**, Profile of yellow line shown in **a** indicating peak signal level and background level. Empty gland lumens indicated by orange arrows in **a** show up as regions of background signal in **b**. **c**, Zoom in of a background region indicated by red box in **b** shows random noise level.

**! CAUTION** Experiments using human tissues must follow appropriate institutional and governmental regulations with respect to informed consent. The results in this figure were from de-identified tissues provided by an institutional tissue bank and are not considered human-subjects research.

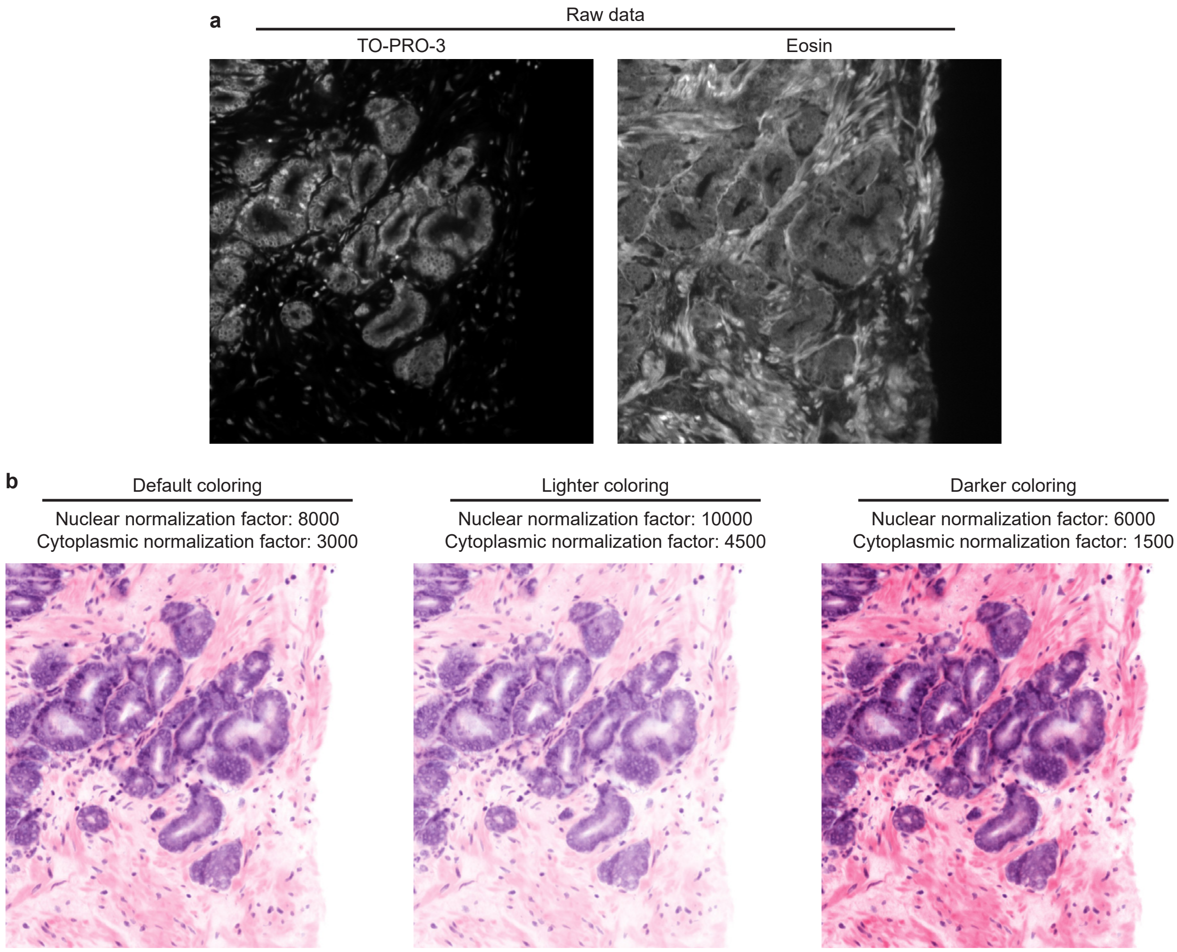


**Fig. S6 | False coloring settings.** The same raw fluorescence data, **a**, can be used to generate false-colored data with a range of appearances, **b**, by adjusting the color parameters (such as normalization factors) in FalseColor Python (step 18).

! CAUTION Experiments using human tissues must follow appropriate institutional and governmental regulations with respect to informed consent. The results in this figure were from de-identified tissues provided by an institutional tissue bank and are not considered human-subjects research.

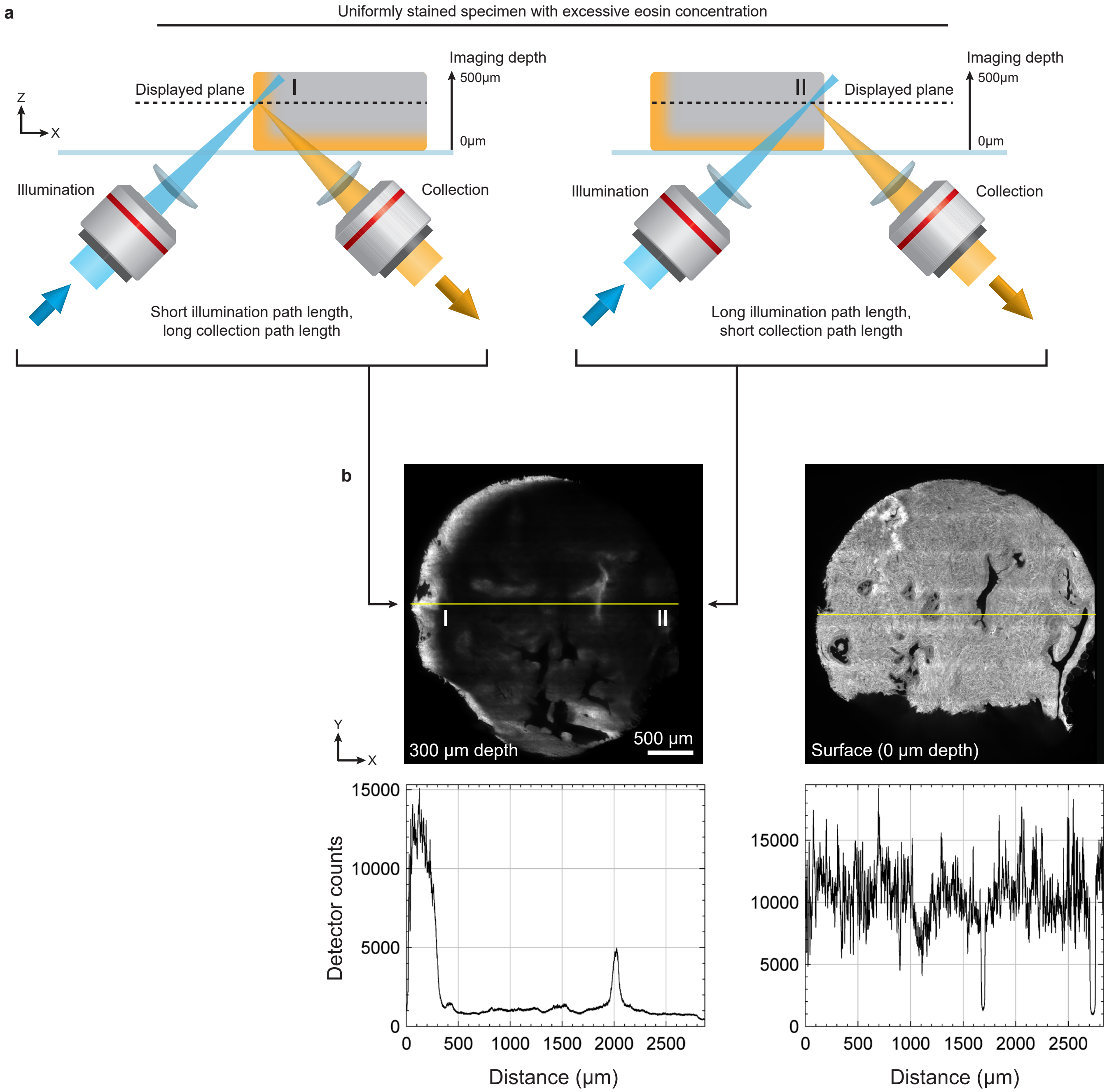


**Fig. S7 | Schematic of beam paths for troubleshooting.** Understanding light paths of a given microscope can be useful to troubleshoot image quality issues. **a,** In this example, a uniformly stained specimen is imaged on an open-top light-sheet (OTLS) microscope. Due to excessive eosin concentration, illumination light is rapidly attenuated by dye absorption at the tissue surface. This results in bright signal at the edges facing the illumination beam (I) and low signal at the tissue core and edges facing away from the illumination beam (II). **b,** Upon initial inspection, *en face* views of the dataset can give the appearance of poor dye penetration, illumination light attenuation, or collection light attenuation. However, evaluation of light paths reveals that high signal is seen in areas of short illumination path length in tissue (I) and that low signal is seen in areas of long illumination path length in tissue (II), suggesting that illumination attenuation due to excessive dye concentration is the culprit.

**! CAUTION** Experiments using human tissues must follow appropriate institutional and governmental regulations with respect to informed consent. The results in this figure were from de-identified tissues provided by an institutional tissue bank and are not considered human-subjects research.

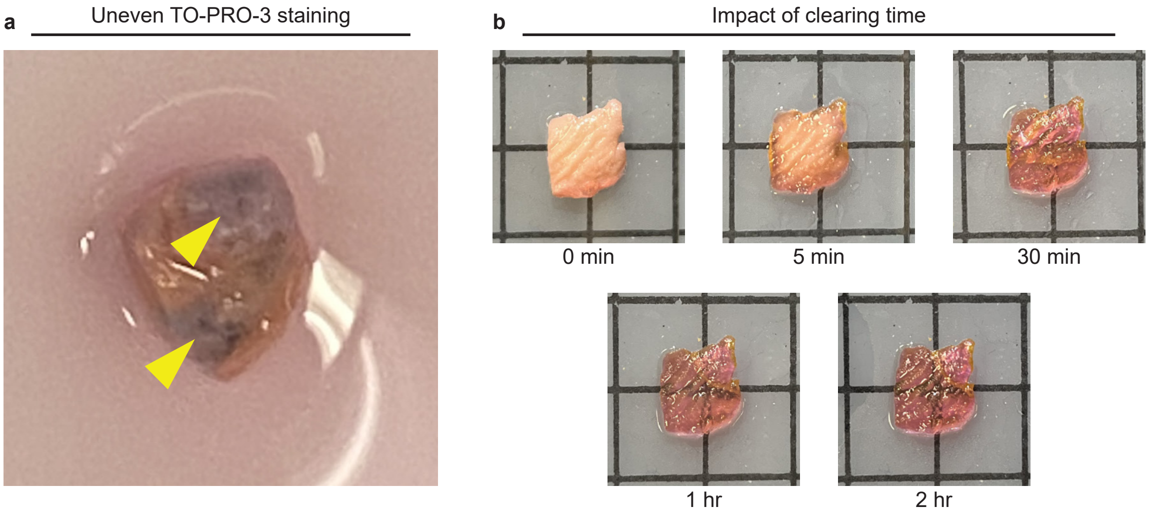


**Fig. S8 | Photos showing visual appearance of tissue after tissue preparation problems. a**, Uneven TO-PRO-3 staining of a human prostate specimen results in visible dark patches on the tissue surface (yellow arrows). **b**, Mouse flank skin specimen shown after increasing amounts of time in ECi clearing medium. Short clearing times result in a relatively opaque specimen. By 2 hours of clearing, the specimen is relatively transparent and suitable for imaging.

**! CAUTION** Experiments using human or animal tissues must follow appropriate institutional and governmental regulations with respect to informed consent/care of animals. The images in panel **a** were from de-identified tissues provided by an institutional tissue bank and are not considered human-subjects research. The tissues shown in panel **b** were obtained from euthanized animals generously provided by veterinarians at an institutional animal facility at the University of Washington. These experiments therefore did not require approval from the University of Washington Institutional Animal Care and Use Committee.

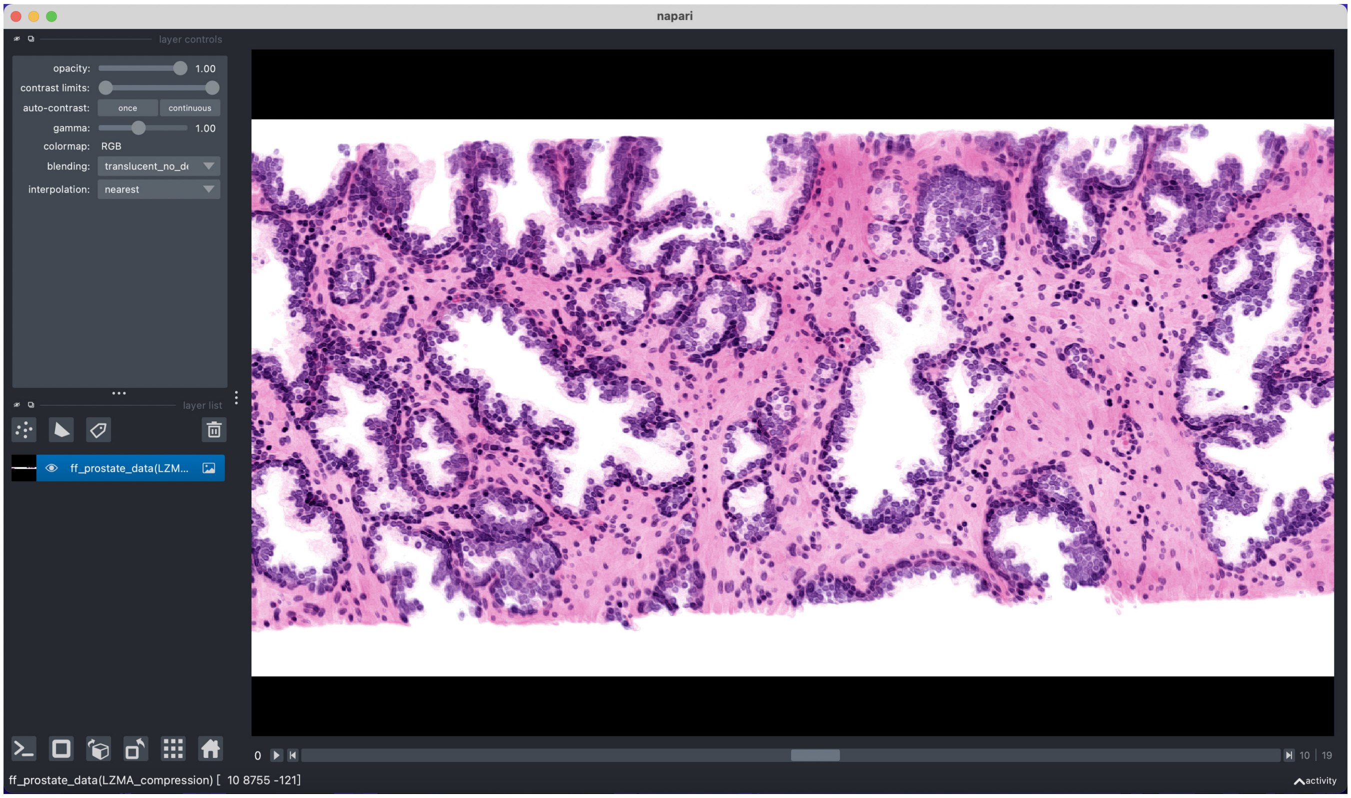


**Fig. S9 | Napari interface for interacting with Path3D data.** Napari viewer (steps 19 - 20), showing a false-colored Path3D dataset of a human prostate core needle specimen. The interface allows users to pan, zoom, and scroll to different depth levels, as well as add annotations.

**! CAUTION** Experiments using human tissues must follow appropriate institutional and governmental regulations with respect to informed consent. The results in this figure were from de-identified tissues provided by an institutional tissue bank and are not considered human-subjects research.

**Fig. S10** Alternative interfaces for interacting with Path3D data. **a**, QuPath interface, showing a false-colored human esophagus biopsy. Pathologist annotations indicating regions of low-grade dysplasia (yellow) and high-grade dysplasia (red). **b**, Custom web interface developed by Ground Truth Labs showing a false-colored human prostate core needle specimen. The integrated grading section below the image allows pathologists to record information such as Gleason grade and presence of cribriform glands for each specimen.

**! CAUTION** Experiments using human tissues must follow appropriate institutional and governmental regulations with respect to informed consent. The results in this figure were from de-identified tissues provided by an institutional tissue bank and are not considered human-subjects research.
